# A split-control criterion for few-shot adaptation in single-cell perturbation prediction

**DOI:** 10.64898/2026.10.05.756906

**Authors:** Wan-Li Liang, Ya-Bing Yang, Qiong-Yi Zhang, Jing-Cheng Feng, Qian-Fei Wang, Dehua Zou, Jia-Lin Tang, Yang Liu, Si-Yuan Wang, Junwei Duan, Yu Bai, Yi-Fang Li, Wan-Yang Sun, Yun-Feng Cao, Rong-Rong He

**Affiliations:** State Key Laboratory of Mechanism and Quality of Chinese Medicine/School of Pharmacy/Faculty of Chinese Medicine, Faculty of Medicine, Macau University of Science and Technology, Macau 999078, China; Shanghai-MOST Key Laboratory of Health and Disease Genomics, NHC Key Lab of Reproduction Regulation, Shanghai Institute for Biomedical and Pharmaceutical Technologies, Shanghai, China; State Key Laboratory of Bioactive Molecules and Druggability Assessment/Guangdong Basic Research Center of Excellence for Natural Bioactive Molecules and Discovery of Innovative Drugs/Guangdong Engineering Research Center of Traditional Chinese Medicine & Disease Susceptibility/Guangdong Engineering Research Center of Traditional Chinese Medicine & Health Products/International Cooperative Laboratory of TCM Modernization and Innovative Drug Development of Chinese Ministry of Education (MOE)/School of Traditional Chinese Medicine/The First Affiliated Hospital of Jinan University/The Second Affiliated Hospital of Jinan University/College of Pharmacy, Jinan University, Guangzhou 510632, China; Institute for Intelligent Bioinnovation, Knowledge Transfer Center, City University of Macau, Macau 999078, China; Ningxia Institute of Medical Sciences, Ningxia Medical University, Yinchuan 750004, PR China; Henry Samueli School of Engineering, University of California, Irvine, Irvine, CA 92697, USA; Faculty of Data Science, City University of Macau, 999078, Macao Special Administrative Region of China; Department of Electrical and Computer Engineering, California State University Fullerton

## Abstract

Few-shot adaptation can help or harm single-cell perturbation prediction, but conventional evaluation shares a control estimate between calibration responses and held-out targets. We show how methods carry control error into their predictions and derive its contribution to score inflation. Nested control-depth experiments support inverse-depth scaling of inner-product inflation; correlation inflation also depends on the control noise-to-signal ratio. Independent-control scoring changed the rank of 40 of 52 configuration–dataset entries. We introduce SafeDelta, a split-control criterion that uses calibration data to decide whether to adapt. Across six datasets, mean gated gain was non-negative under the original two-control evaluation. On a deep plate, fixed predictions and decisions retained positive mean gains under three additional independent control references. The same reproducibility principle identified a fibrosis programme that localises to scar in human liver. These results inform adaptation decisions and control-depth planning.

---

Models that predict single-cell transcriptional responses to genetic and chemical perturbations^1,2^ are increasingly asked to work in contexts they were not trained on: a new cell line, a new patient sample, a new tissue. The space of interventions is combinatorially large, the number that can be assayed in any one context is not, and a model that transports would let responses measured in one context stand in for the same perturbations in another^3–7^. Existing methods transport perturbation information across contexts using latent representations, compositional models or distributional mappings^2,8–13^. Fully characterising a new context is expensive, motivating the few-shot setting considered here: a handful of responses are measured in the new context and used to correct a shared predictor, and the correction is then applied to unmeasured perturbations.

The compromise inherits a question that has to be answered before the remaining responses are measured. A correction estimated from a handful of noisy responses can degrade a prediction rather than improve it, and the quantity that would tell the two cases apart is the true gain on perturbations not yet measured, which is precisely what is being predicted. Nor can a practitioner fall back on a published ranking, because those rankings are computed under the same control convention^14–16^, and published evaluations show that conclusions about model performance depend on the evaluation protocol^17–19^.

The reason lies in how these experiments are scored. Every perturbation in a context is referenced to a single control estimate, and a recent analysis showed that this shared control inflates apparent performance, can distort the relative ranking of algorithms, and is removed by scoring against an independent half of the controls^20^; it is an instance of the leakage between what is fitted and what is scored that is a recognised cause of over-optimistic performance^21,22^. That analysis conditions on the prediction, treating it as a deterministic vector, so the control estimate enters the score only through the metric that subtracts it, and the coefficient with which the control’s error is carried into the score is fixed at one. It also gives the dependence of the bias on control depth, and shows that the split-control estimator is conservative, attenuated towards zero. Few-shot adaptation violates the conditioning assumption. The correction is built from measured responses in the target context, each referenced to the same control estimate the evaluation will use, so the control’s error is carried into the prediction itself and not only into the metric. Here we extend the analysis to predictors that read the target context’s controls. We show that the coefficient is a property of a method together with the pipeline it is run in, that it can be estimated by rebuilding the prediction from each control half and projecting their difference, and that it takes exact values for whole classes of estimators: zero for predictors that never read the target context’s control, and one when the mean calibration response is added at unit weight to a control-independent predictor.

Under the carriage model, inner-product inflation is inverse in control depth, while PCC inflation also depends on the control’s noise-to-signal ratio, which we test in a same-plate nested control-depth experiment. The law can be read in two directions. Inverted, it converts a question about bias into a question about experimental design: what control depth is associated with a specified scoring-bias target. Read forward, it yields a criterion, computable from calibration data alone, that decides whether to adapt in a given context; abstaining returns the shared predictor unchanged, so a decision not to adapt costs nothing that was already working. The criterion guarded against harmful adaptation: where controls were shallow it abstained throughout and avoided the loss, and where they were deep its realised gain was marginally below that of adapting every time. This reinforces the importance of benchmark design^23–25^: a leaderboard whose reference measurement is shared within a context reports a quantity that is partly a property of the evaluation. Finally, the principle behind the criterion, that what reproduces across independent estimates is what transports, is not specific to model evaluation: applied to a fibrosis programme defined in vitro, it selects the part that localises to scar in human liver^26–28^.

## Results

### Calibration data guide few-shot adaptation

We considered a minimal adaptation interface in which a frozen shared predictor, here the mean response to the same perturbation in the other contexts, provides a baseline response for every perturbation in the target context, and adaptation adds a context-specific correction that is estimated from a small calibration set of measured responses (Fig. 1a). Figures and tables call this frozen predictor Pooled. Withholding the correction recovers the frozen predictor exactly. The question is whether a correction estimated from k calibration responses transfers to perturbations that were not measured.

**Figure 1.**
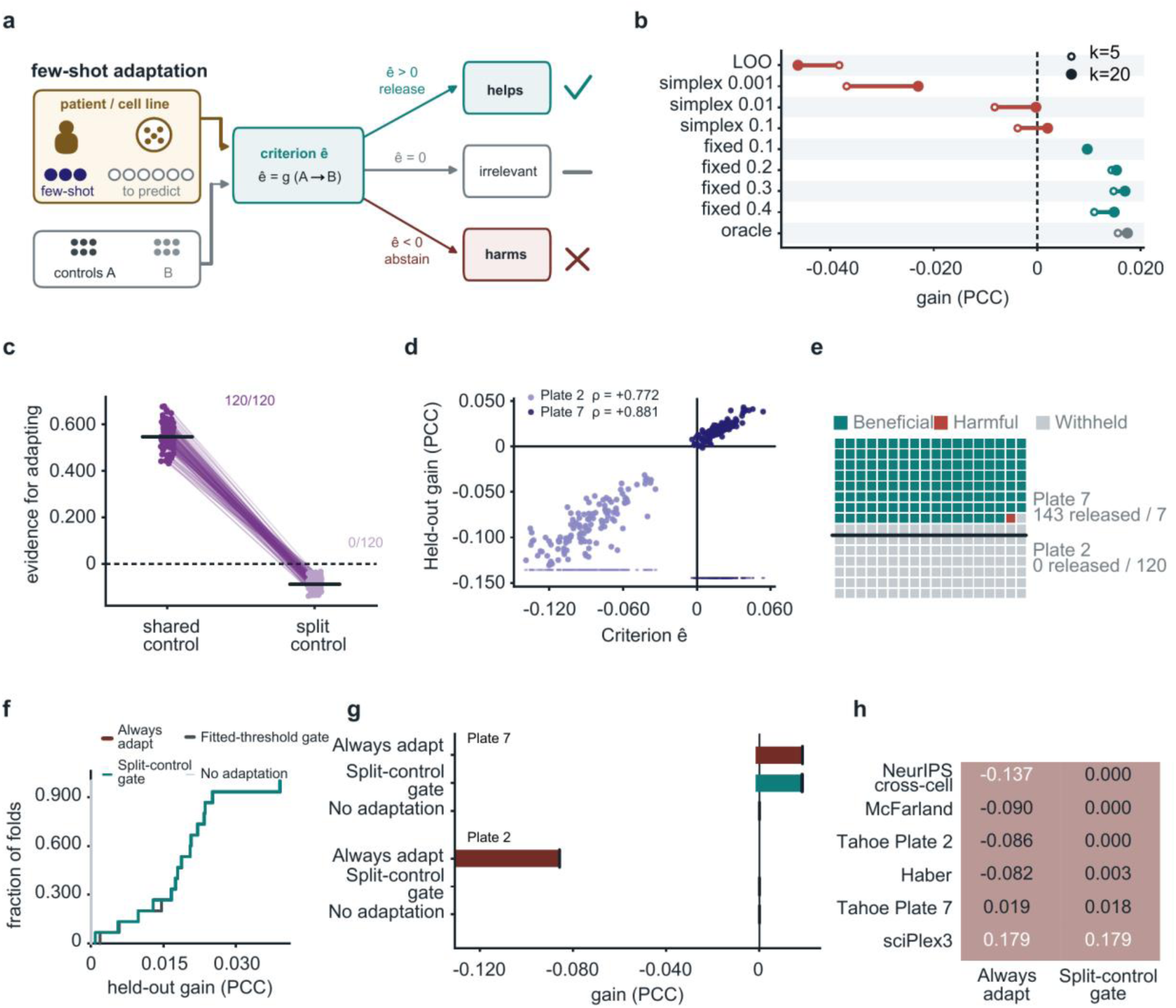
A split-control criterion guides few-shot adaptation using calibration data alone. **a**, Few-shot adaptation. A new context carries *k* measured responses and many unmeasured ones; its control cells are split into disjoint halves A and B. The criterion *e* (equation (10)) is the leave-one-out gain of the correction on the calibration responses, built against half A and scored against half B; the correction is released when *e* > 0 and withheld otherwise. **b**, Held-out gain over the shared predictor for adaptation weights chosen by leave-one-out (LOO), by simplex fitting at three ridge strengths, by fixed shrinkage (0.1 to 0.4) and by an oracle, at *k* = 5 (open) and *k* = 20 (filled); deep plate, 16 cell lines, 20 calibration draws per line. **c**, Evidence for adapting computed against the shared control and against the split control, one line per trial (shallow plate, *n* = 120 trials); black bars, medians. **d**, Criterion against observed held-out gain, one point per trial (shallow plate, *n* = 120; deep plate, *n* = 150). **e**, Release decision, one square per trial, on the deep plate (*n* = 150) and the shallow plate (*n* = 120): released and beneficial, released and harmful, or withheld. **f**, Cumulative distribution over 15 leave-one-context-out folds of the held-out gain on the deep plate under four policies: always adapt, Fitted-threshold gate, Split-control gate, and never adapt. **g**, Gain for three policies on both plates; bars span the worst-trial gain to the mean, and black ticks mark the mean. **h**, Mean held-out gain under unconditional and gated adaptation on six datasets, using disjoint control halves. The atlas plates use a mixture adapter and the external datasets an offset adapter, with each exchanged A/B direction scored separately before averaging. Throughout, gain is the change in Pearson correlation across genes relative to the shared predictor. In c-e and g, each trial is one target context evaluated for one calibration draw.

The evidence conventionally used to answer this question favours adapting whether or not adapting helps. The apparent gain of the correction on the calibration responses is computed against the same control estimate from which the correction is derived, so the correction is rewarded for reproducing the sampling error of the control, and this spurious gain increases with control noise. To quantify the effect, we analysed two plates of a drug-response atlas^29^, one with 10 and one with 400 control cells per half (the shallow and deep plates), defining a trial as one target context scored for one calibration draw. In a separate weight comparison on the deep plate, fitting the adaptation weight on the conventional evidence reduced accuracy relative to no adaptation (Fig. 1b and Extended Data Fig. 1a, b), and adapters of higher capacity performed no better (Extended Data Fig. 1c, d). On the shallow plate, the conventional evidence was positive in all 120 trials (Fig. 1c), whereas the observed held-out gain was negative in all 120 (Fig. 1d). On this plate, the conventional evidence recommended adapting in every case in which adapting was harmful.

To remove the shared term, we devised a split-control criterion. The control cells of the target context are divided into two disjoint halves; the correction is estimated against one half and evaluated against responses referenced to the other, so that agreement arising from a common control estimate is removed by construction. The criterion requires no held-out perturbation and can be computed as soon as the calibration responses are available. Within each plate it tracked the observed held-out gain (Spearman *ρ_s_* = +0.772 on the shallow plate and +0.881 on the deep plate; Fig. 1d). We released the correction when the criterion was positive and withheld it otherwise, without a tuned threshold. In this evaluation, half B supplied both the decision evidence and the held-out scoring. No trial was released on the shallow plate, which avoided the mean loss of 0.0858 incurred by unconditional adaptation. On the deep plate, where unconditional adaptation yielded a mean gain of 0.0185, 143 of 150 trials were released, of which 142 had a positive observed gain, and 7 of the 149 beneficial trials were withheld (Fig. 1e). The criterion avoided the shallow-plate loss while retaining most of the deep-plate gain.

This protection came at a small cost. On the deep plate the criterion lowered the mean gain by 0.0002 relative to unconditional adaptation, because it withheld seven beneficial trials and released the single harmful one. A threshold tuned by leave-one-context-out validation recovered this difference by releasing every trial (Fig. 1f); transferred to the shallow plate, it released nine trials, all of which were harmful. On the shallow plate the criterion avoided the mean loss of 0.0858 and a worst-trial loss of 0.1308 under unconditional adaptation (Fig. 1g).

With predictions and release decisions held fixed, three additional disjoint control references preserved positive mean gated gains on the deep plate (0.0177–0.0189; all 95% context-bootstrap intervals above zero; Extended Data Fig. 2a). Evidence–gain correlations remained positive but decreased from 0.88 to 0.58–0.69 (Extended Data Fig. 2b). Of the 143 released trials, 0–3 had negative observed gains under each additional reference, with none negative under all three (Extended Data Fig. 2c). Four released trials changed gain sign across references (Extended Data Fig. 2d).

We next asked whether the rule generalises beyond the plates on which it was developed. We applied the same zero-threshold rule to four further datasets spanning two assay families, splitting each dataset’s own control cells by a key independent of the perturbation (Extended Data Fig. 3a, b): a chemical screen^30^, an intestinal epithelial atlas^31^, a multiplexed cancer cell-line screen^32^ and the NeurIPS cross-cell benchmark^15,33^. The atlas plates used a mixture adapter; these external evaluations used an offset adapter, with each exchanged A/B direction rebuilt and scored before the two were averaged. Unconditional adaptation was beneficial in one dataset and harmful in three, with mean gains ranging from +0.179 to −0.137, and the conventional shared-control evidence released 76 to 100% of the scored responses in the harmful datasets. Under the criterion, every scored response was released in the chemical screen and none in the two most harmful datasets. Under the original two-control evaluation, mean realised gain was non-negative in all four external datasets and in both atlas plates, six of six datasets in total (Fig. 1h). The rule carried over to datasets that played no part in its development, although the near-uniform release or abstention within several datasets indicates that much of the observed discrimination was between datasets.

### Shared-control inflation exceeds the differences the benchmark reports

A simpler alternative to a per-context decision would be to adopt the top-ranked method of a published benchmark^14–16^, so we asked whether such rankings can be resolved. We rescored three generative models (trVAE^10^, scGen^9^ and scPRAM^12^) on the four datasets of a published leaderboard^15^ under independent controls, splitting each dataset’s controls by a key independent of the perturbation (Extended Data Fig. 3a, b), and estimated the difference between each pair of adjacent ranks with 95% intervals obtained by bootstrapping whole contexts. Two of the 8 adjacent pairs were separated from zero, and the top-ranked pair was not separated in any of the four datasets (Fig. 2a). Under the benchmark’s own shared-control scoring, 3 of the 8 pairs were separated. The leaderboard does not identify a best method on any of these datasets.

**Figure 2.**
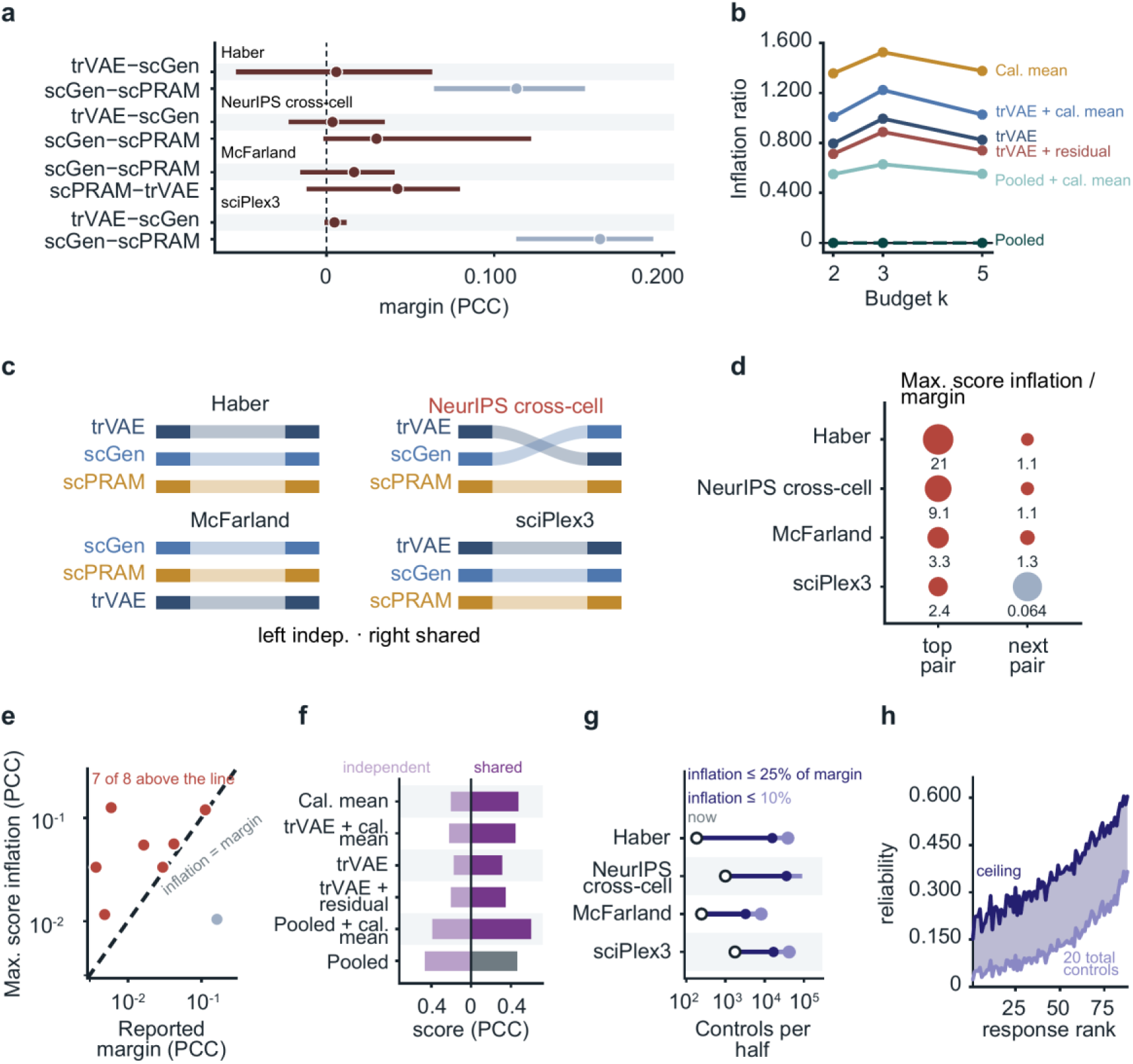
Shared-control inflation exceeds the margins a published benchmark reports. **a**, Paired context-perturbation difference in Pearson correlation (PCC) between adjacent ranks of a published leaderboard^15^ under independent controls, for three generative models on four datasets (8 pairs). Points, mean; bars, 95% interval from a bootstrap of whole contexts (2,000 resamples). **b**, Relative inflation (shared minus independent score, divided by the independent score) against calibration budget *k* for six prediction configurations on six cell lines of the deep plate (five at *k* = 3); controls were split into halves of 250 cells (198 and 199 in one line). Pooled + cal. mean adds the mean calibration response to Pooled at weight one. **c**, Method ordering per dataset under independent (left) and shared (right) controls. **d**, Ratio of the larger of each pair’s two individual-score inflations to the reported margin, for the top and the next adjacent pair of each dataset; bubble size indicates departure from 1 on a logarithmic scale; colour distinguishes ratios above and below 1. **e**, Maximum score inflation against margin for the 8 pairs on log axes; dashed line, inflation equal to margin. **f**, PCC at *k* = 2 under shared and independent controls for the methods in **b**. **g**, Projected control depth per half for reducing inflation below 25% (filled) and 10% (open) of the smallest adjacent margin, extrapolated from the present benchmark-subset depth (circle) using the observed PCC inflation, with the denominator of equation (7) held constant. **h**, For the 88 drugs of the shallow plate, response reliability at 20 control cells in total and its implied correlation ceiling (the square root of each drug’s reliability), ordered by response magnitude.

Sharing controls shifted scores by more than the differences between methods. We define the inflation of a score as its value under shared controls minus its value under independent controls. Inflation did not diminish as more calibration responses were measured: for six prediction configurations compared on six cell lines of the deep plate, the relative inflation of each of the five configurations that read the target context’s controls was at or above its k = 2 value at k = 5, and that of the shared predictor, which never reads them, was zero at every calibration budget (Fig. 2b). In the leaderboard, rescoring under independent controls preserved the order obtained under shared controls in three of the four datasets and exchanged the top two methods in the fourth (Fig. 2c). Taking the larger of each pair’s two individual-score inflations, the ratio of inflation to margin reached 21 and had a median of 2.9 across the six unresolved pairs; the pair with the smallest margin (0.0037) carried an inflation 9 times that margin (Fig. 2d). The inflation exceeded the margin in 7 of the 8 pairs (Fig. 2e). In most of these comparisons, the quantity being ranked is smaller than the error with which it is measured.

The size of the inflation provides a projection of the control depth needed to limit scoring bias. Methods that read the target controls scored 1.5 to 2.4 times higher under shared than under independent controls, whereas the shared predictor scored identically under both (Fig. 2f). With the PCC denominator held constant, inverse-depth extrapolation placed the depth needed to reduce inflation below a quarter of the adjacent margin at roughly 10 to 85 times the present depth per control half across the four datasets, from about 10-fold for the deepest benchmark subset (1,752 control cells per half) to about 85-fold for the shallowest (184); at a tenth of the margin the projected multiplier was 24 to 213 (Fig. 2g).

Measurement reliability also limits the interpretation of a ranking. On the shallow plate, split-half response reliability averaged 0.140 over 88 drugs at 20 control cells in total; the corresponding mean of the per-drug square-root ceiling estimates was 0.357 under an independent measurement-error model (Fig. 2h).

### Inflation scales with the control error that a method carries

We then asked how published methods compare once the controls are independent. We scored thirteen prediction configurations on four public datasets (Extended Data Fig. 3a), 52 configuration-by-dataset entries, under shared and under independent controls, with each configuration’s predictions and treated profiles held fixed between the two scoring regimes. Under independent controls, the pooled baseline, which never reads the target context’s controls, ranked first in three of the four datasets (0.706, 0.598 and 0.563) and was outscored in the chemical screen, where it ranked twelfth of the thirteen configurations at 0.283 and the calibration mean led at 0.493 (Fig. 3a). The methods that lost ground were the ones that read the target controls.

**Figure 3.**
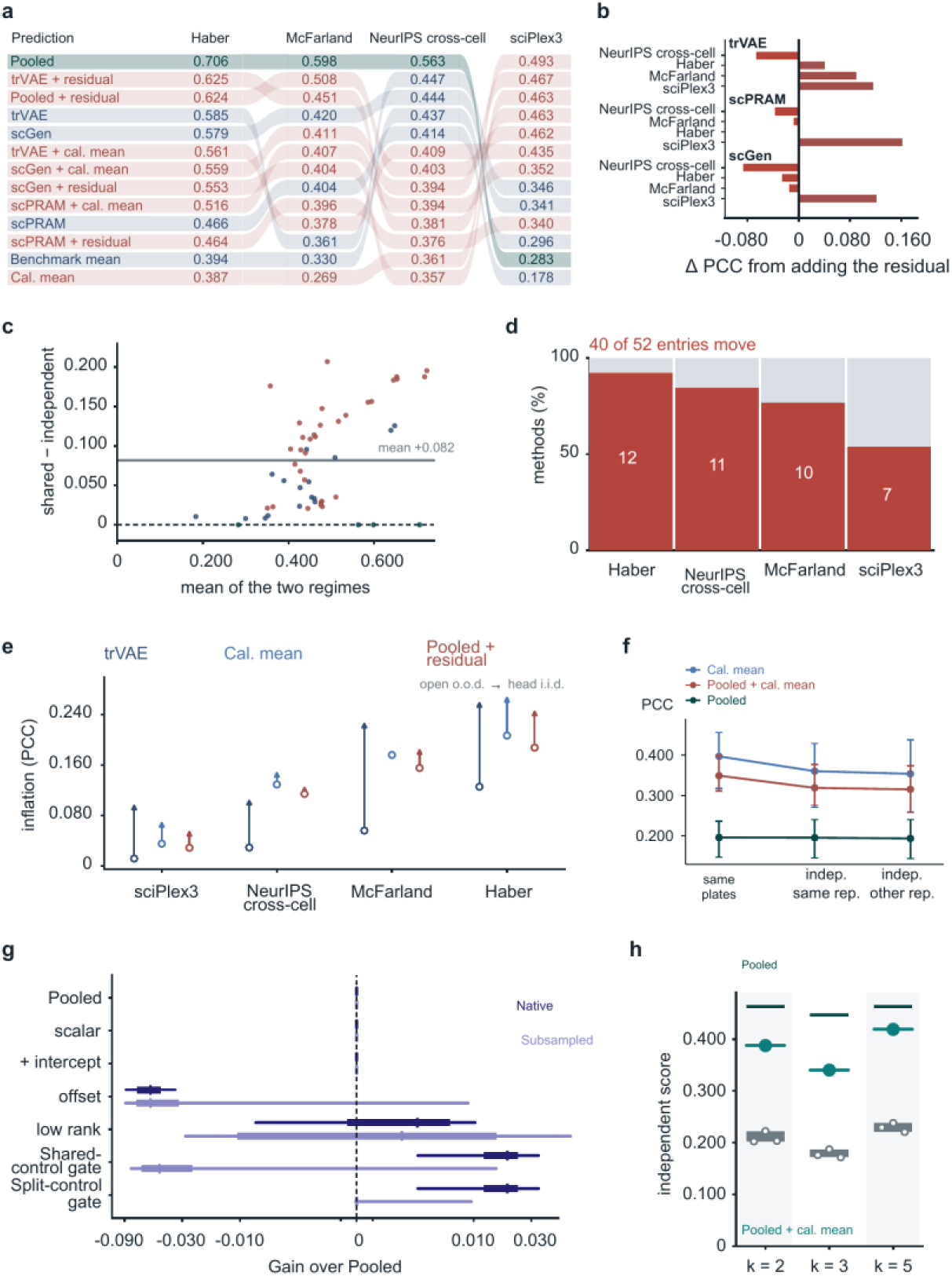
Prediction configurations under independent controls, and how much control error each carries. Thirteen prediction configurations on four public datasets (52 configuration-by-dataset entries), each scored under shared and under independent controls holding predictions and treated profiles fixed between regimes. **a**, PCC under independent controls for thirteen configurations. Equal-width ribbons connect the same configuration across datasets; row names follow the Haber order, and each dataset column is ordered on its own unrounded scores, which are printed to three decimals. Colour distinguishes the control-free pooled baseline, published-model and benchmark-mean configurations, and additive forms. **b**, Change in independent-control PCC when a context-specific residual is added to trVAE, scPRAM or scGen. **c**, Inflation (shared minus independent PCC) against the mean of the two scores for the 52 entries; grey line, mean inflation over all 52 entries. **d**, Share of entries whose rank changes between regimes, per dataset; column width proportional to entries per dataset. **e**, Inflation under the out-of-distribution split (open) and an i.i.d. split (arrowhead) for three estimators. **f**, PCC of Cal. mean, Pooled + cal. mean and Pooled, scored against controls shared with the prediction, an independent split of the same replicate, or controls from another replicate; means and 95% hierarchical bootstrap intervals over three contexts (*n* = 471 scored responses per regime). **g**, Gain over Pooled in independent-control PCC for each adapter at native depth (dark; 400 treated cells per condition and 400 controls per half) and subsampled depth (light; 20 treated cells and ten controls per half); 15 cell lines, 12 calibration draws each (*n* = 180 trials per depth). Boxes, median and interquartile range; whiskers, 5th to 95th percentiles; symmetric-log axis, linear within ±0.005. **h**, Independent-control PCC at *k* = 2, 3 and 5 on six cell lines of the deep plate (five at *k* = 3) for Pooled + cal. mean at weight one (filled), three other adaptation schemes (open) and the control-free predictor (line).

How much a method loses is set by how much of the control error it carries. We define the carriage coefficient of a method as the amount of the control estimate’s error that its prediction carries, in units of that error. The coefficient is zero for a method that never reads the target context’s controls and one when the mean calibration response is added at unit weight to a control-independent predictor. Estimated by projection, published models run on their own lay between these two values in every dataset we measured (0.267 to 0.810), and one model varied across datasets as much as different models varied from each other (Extended Data Fig. 4a). A predictor that adds a calibration mean to such a model carries the sum of the two coefficients, which exceeds one (Extended Data Fig. 4b). On the deep plate, the coefficient was exactly 0 for the shared predictor, exactly 1 for the calibration mean and 0.96 for a model fine-tuned on the target controls (Extended Data Fig. 4c). The norm ratio overestimated the projection coefficient for configurations built on a published model (Extended Data Fig. 4d), and within each dataset the published model run on its own had the smallest coefficient and the smallest inflation (Extended Data Fig. 4e, f). The coefficient is a property of a method together with the pipeline and dataset it is run on.

Inflation was confined to the entries that read the target controls. Attaching a context-specific residual to a published model did not improve it consistently: across the four datasets the independent-control score changed by +0.045 for trVAE, +0.029 for scPRAM and −0.002 for scGen, the largest gain for each model came from the chemical screen (+0.116, +0.162 and +0.121), and the pooled effect was not separated from zero (Fig. 3b and Extended Data Fig. 3c). Across all 52 configuration-by-dataset entries, inflation averaged +0.082 and increased with the score being inflated; it was zero for the four control-free entries (Fig. 3c). The methods that look best under shared controls are the ones whose scores are inflated most.

Making the controls independent changed the ranking. Of the 52 entries, 40 changed rank, with changes in all four datasets (12, 11, 10 and 7 entries; Fig. 3d). The inflation appeared under both ways of holding perturbations out: with a random split in place of the out-of-distribution split it remained between 0.057 and 0.270 for the three estimators tested, and the top-ranked estimator changed only in the intestinal atlas (Fig. 3e). For the published models, the shared-control score exceeded the independent-control score in every context: by 0.23 to 0.37 in the 16 cell lines scored with scGen and by 0.12 to 0.19 in the six scored with trVAE on the deep plate, and by 0.005 to 0.017 in the three contexts of the chemical screen scored with trVAE (Extended Data Fig. 3d-f). The inflation also appeared within a single experiment. When the scoring controls were an independent split of the same replicate, the calibration mean fell from 0.397 to 0.361 and the shared predictor plus calibration mean from 0.350 to 0.319; with scoring controls from a separate replicate the two scores were 0.354 and 0.316. The control-free predictor was unchanged across the three scoring controls (0.196, 0.196 and 0.194; Fig. 3f). A published ranking reflects the scoring convention as much as it reflects the methods.

For our own adapters, accuracy depended on the decision to adapt and not on adapter capacity. We compared adapters of increasing capacity on the deep plate under independent controls, at native depth (400 treated cells per condition and 400 controls per half) and after downsampling to 20 treated cells and ten controls per half (Fig. 3g and Extended Data Fig. 5a, b). Relative to the shared predictor, a global offset lowered the score at both depths (−0.0582 and −0.0492), a low-rank correction changed it by +0.0026 and +0.0048, and a fitted mixture of these corrections, released by the shared-control gate, gained 0.0183 over the 180 trials of this comparison at native depth but lost 0.0418 at subsampled depth. Released by the split-control gate, the same mixture kept the gain at native depth and removed the loss at subsampled depth (+0.0007). With an oracle weight, eight calibration responses gave 90% of the gain obtained with 76 (Extended Data Fig. 5c). In the separate budget comparison, the shared predictor plus the calibration mean, applied at full weight without shrinkage or gating, was the strongest of the methods that read calibration responses at every budget (0.388, 0.340 and 0.419 against 0.172 to 0.238), yet at every budget it scored below the control-free predictor (0.463, 0.447 and 0.463; Fig. 3h and Extended Data Fig. 5d). The criterion also ran unchanged on a published conditional generative model^10^ used as the frozen base: its cost relative to unconditional adaptation was at most 0.019 on the three external datasets in which adaptation helped, precision among the released scored responses was 0.67 to 0.88, and mean gated gain was non-negative in all four (Extended Data Fig. 6a, b). Our re-runs reproduced the published scores of six method-by-dataset pairs to within 0.026 (Extended Data Fig. 6c), and the top two configurations exchanged rank only where the difference between their inflations exceeded their shared-control margin (Extended Data Fig. 6d).

### Inflation tracks the control noise-to-signal ratio across settings

The size of the inflation is set by the control’s noise-to-signal ratio. To screen this quantity without a model, we computed, for 99 context-by-split estimates in eight dataset partitions, half the centred squared distance between the control half-means divided by the uncentred response inner product. This screening ratio ranged from 0.0008 to 2.454, a 3,101-fold span, and exceeded 0.1 in 42% of the estimates (Fig. 4a); the theoretical ratio *λ* of equation (1) uses the centred response signal in its denominator instead. Both components varied substantially: the response signal ranged from 5.55 to 364.18 (approximately 66-fold), and control noise from 0.175 to 30.40 (approximately 174-fold; Fig. 4b). The screening ratio therefore reflects variation in both control noise and response strength.

**Figure 4.**
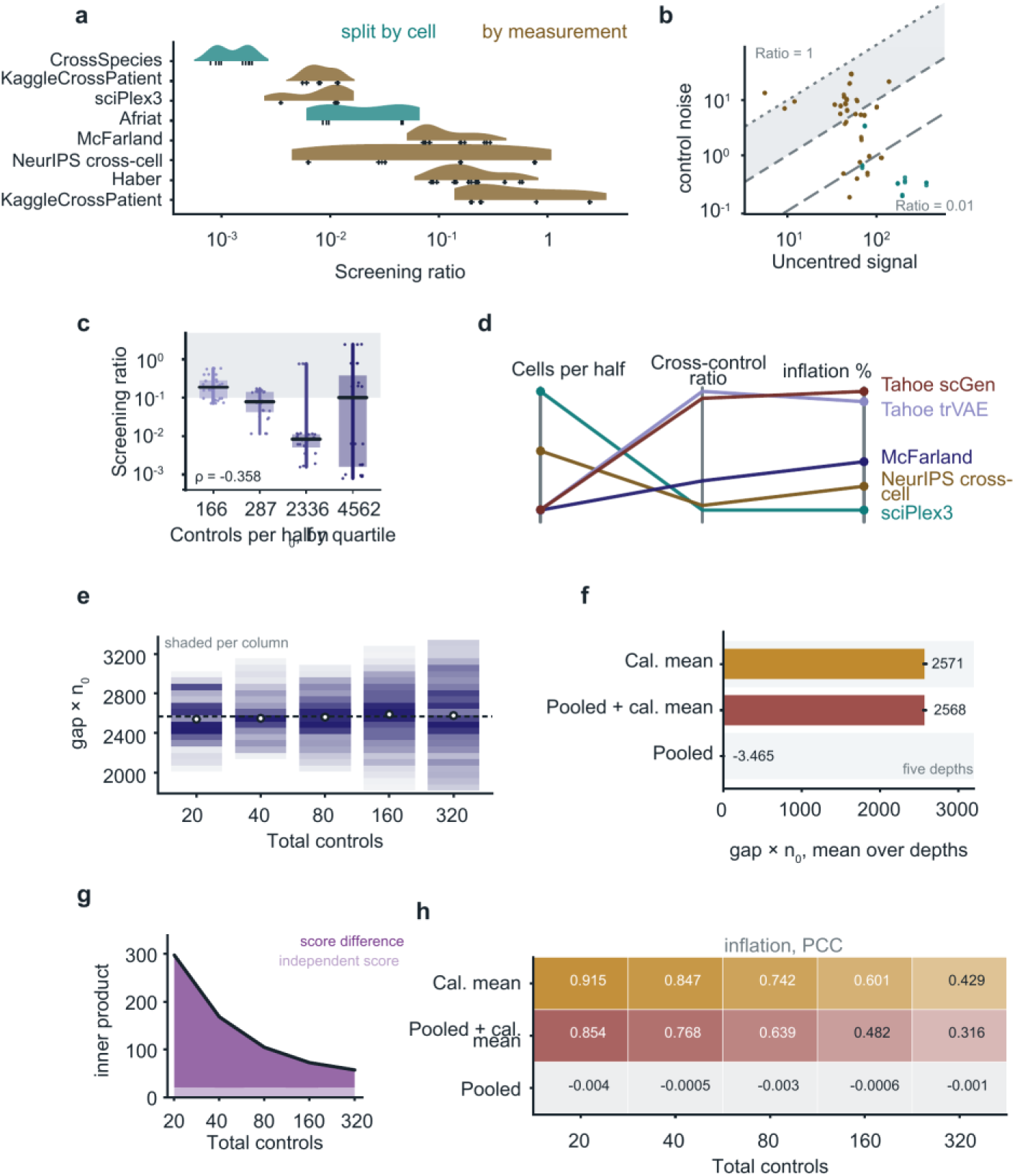
Shared-control inflation depends on the noise-to-signal ratio and control depth. **a**, Screening ratio for 99 context-by-split estimates across eight dataset partitions: half the centred squared distance between the control half-means, divided by the uncentred response inner product; colour, controls split at the cell level (2 partitions) or by a measurement key (6 partitions). **b**, The same estimates as control-error energy against the uncentred response signal; dashed lines, screening ratios of 1 and 0.01. **c**, Screening ratio by quartile of control depth per half (median depth per quartile on the axis); boxes, interquartile range with median; whiskers, minimum to maximum; grey band, screening ratio > 0.1; *ρs*, Spearman correlation between depth and screening ratio. **d**, Control depth per half, the cross-control ratio and the inflation of the calibration-mean score for five settings: three datasets scored by the published benchmark pipeline, and the deep plate scored with scGen and with trVAE; each axis scaled to its own range. **e**, Product of the inner-product gap and control depth per half *n*_0_ (equation (11)) for Pooled + cal. mean, per scored response as the deep-plate control halves are subsampled from 10 to 160 cells (20-320 in total; *n* = 800 scored responses per depth); shading, column-wise density; circles, medians; dashed line, pooled mean. **f**, The same product averaged over the five depths for three estimators; black lines, range across depths. **g**, Shared-control inner product decomposed into the independent-control inner product and their difference, against total control cells. **h**, Inflation on the PCC scale for the three estimators at each depth.

Control depth alone did not predict the ratio. Across quartiles of control depth (medians of 166, 287, 2,336 and 4,562 cells per half), the median ratio was 0.188, 0.078, 0.008 and 0.101, so that the deepest quartile had the second-highest median. The two deepest quartiles spanned 2.7 and 3.5 orders of magnitude, against 0.9 for the shallowest; 12 of the 24 estimates in the deepest quartile exceeded the median of the shallowest, and depth and ratio were only weakly correlated (Spearman *ρ_s_* = −0.36; Fig. 4c). Across five settings whose control depth differed 7.2-fold, the ratio differed 14.2-fold and the inflation 21.0-fold, and two settings with nearly identical depth (245 and 250 control cells per half) differed threefold in ratio and more than twofold in inflation (Fig. 4d). Deep controls do not guarantee a small inflation.

What remained approximately constant was the product of control depth per half and the inflation of the unnormalised score, the inner product that forms the numerator of the correlation. When the control halves of the deep plate were subsampled in nested steps from 10 to 160 cells (20 to 320 controls in total), the median product over the 800 scored responses at each depth stayed within 1.1% of its pooled value (2,540 to 2,590 against 2,568), while its interquartile width increased from 261 to 522 (Fig. 4e). The constant was specific to the estimator: 2,571 for the calibration mean and 2,568 for the shared predictor plus calibration mean, each with a coefficient of variation near 0.005 across depths, and −3.5 for the control-free predictor (Fig. 4f). For a given estimator and context, the inner-product inflation followed an approximately inverse-depth relationship.

We decomposed the shared-control inner product into the independent-control inner product and the difference between the two. Across the depth ladder, the difference fell by 94% (255.6 to 16.1), while the independent-control inner product fell by 2%; the difference accounted for 92% of the shared-control score at 20 total control cells and 44% at 320 (Fig. 4g). On the correlation scale the relationship was monotonic but not proportional to reciprocal depth: a fit through the origin gave negative coefficients of determination, −1.24 for the shared predictor plus calibration mean and −3.37 for the calibration mean (Fig. 4h).

### Measurement depth shapes evaluation and adaptation

To test the depth dependence directly, we varied depth within one plate. On the deep plate, treated cells were subsampled from 400 to 20 per condition and controls from 400 to ten per half while the cell lines, the perturbations and the calibration draws were left unchanged, and six adapters were scored under shared and under independent controls at both depths (180 trials per depth and scoring convention). The inflation appeared where the law predicts. For the adapter that adds a global offset estimated from the target context, the shared-control score exceeded the independent-control score by a median of +0.224 at native depth and +0.722 at subsampled depth, whereas for the other five adapters the difference was at most 0.092 (Fig. 5a). Absolute scores fell from about 0.67 to about 0.11 at the lower depth, which reflects the loss of measurement reliability.

**Figure 5.**
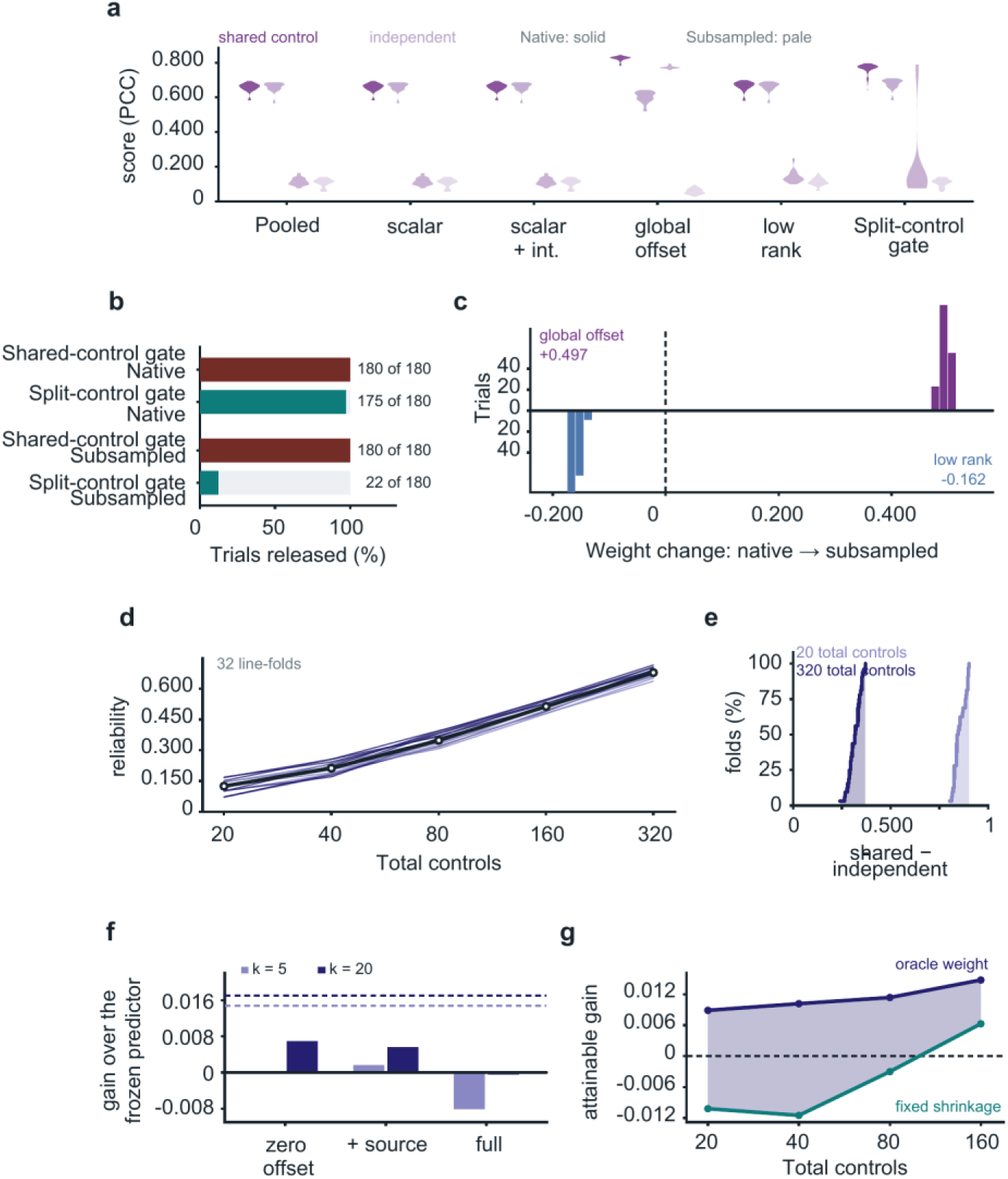
Measurement depth shapes evaluation and adaptation. All panels use the deep plate. **a-c**, Treated cells subsampled from 400 to 20 per condition and controls from 400 to ten per half; 15 cell lines, 12 calibration draws each (*n* = 180 trials per depth and regime). Depth labels 400 and 20 denote treated cells per condition. **a**, PCC of six adapters under shared (dark purple) and independent (light purple) controls at native (solid) and subsampled (pale) depths; violins, distribution over trials. The gated rung uses the split-control criterion. **b**, Share of trials in which the correction is released by the shared-control gate and by the split-control gate at each depth. **c**, Per-trial change in the fitted mixture weight of the global offset (above the axis) and of the low-rank offset (below) when depth is reduced from 400 to 20 cells. **d-e**, Total control cells subsampled from 20 to 320 (10–160 per half), with 150 treated cells per condition fixed; 16 cell lines, 30 drugs, both control-split directions (*n* = 32 line-folds). **d**, Split-half reliability of the measured response against control depth, one line per line-fold; heavy line, median. **e**, Cumulative distribution over the 32 line-folds of the shared-minus-independent score difference at 20 and 320 control cells. **f**, Held-out gain over the frozen predictor at *k* = 5 and *k* = 20 for three adapter forms (zero-mean offset, offset with source term, full adapter), with mixture weights fitted to the calibration responses on the simplex under a ridge penalty of 0.01 towards equal weights; dashed lines, fixed shrinkage of 0.3. Native control depth, independent-control scoring, 16 cell lines and 20 calibration draws per line. **g**, Gain attainable with a correction constructed from all held-out responses, under fixed shrinkage of 0.3 or an oracle weight, against total control cells (20–160; see Methods).

The scoring convention also determined the release decision. On the conventional evidence, the correction was released in all 180 trials at both depths. The split-control criterion released 175 of 180 trials at native depth and 22 of 180 at 20 control cells (Fig. 5b). Of those 22, 13 were beneficial and 9 harmful, and 152 of the 158 withheld trials would have been harmed by adaptation. On the shallow plate, which shares only some of its cell lines and drugs with the deep plate, no trial was released at the same depth. On that plate, a single additive term common to all 88 drugs of a context carried a median of 48% of the response variance (Extended Data Fig. 6e). Removing this term from prediction and truth alike reduced the mean gain of a ridge release policy whose fits were fixed before held-out outcomes were read, labelled Ridge policy (Plate 2) in the figures, from 0.355 to 0.0001 (Extended Data Fig. 6f), and reduced the mean score of the calibration mean from 0.590 to −0.021, the value obtained by centring alone on independent noise (Extended Data Fig. 6g). Fitted parameters shifted in the same way. At the lower depth, the weight of the full offset in the fitted mixture increased in all 180 trials (median +0.497), whereas the weight of the low-rank offset decreased in all 180 (median −0.162; Fig. 5c). Under shared controls, the fitted strength of adaptation follows the depth of the measurement.

With treated-cell profiles held fixed, increasing control depth produced the same pattern. Across a nested ladder from 20 to 320 total control cells (10–160 per half), with 150 treated cells per condition held fixed, the split-half reliability of the measured response rose from a median of 0.125 to 0.678, and at 20 cells it ranged from 0.071 to 0.169 across the 32 cell-line folds (Fig. 5d). The difference between the shared-control and independent-control scores was positive in every fold at both ends of the ladder, with minima of +0.802 at 20 cells and +0.240 at 320 and medians of 0.844 and 0.319 (Fig. 5e).

Additional adapter capacity did not compensate for shallow controls. Relative to the frozen predictor, the gain at k = 5 and k = 20 was 0.0002 and 0.0069 for a zero-mean offset, 0.0017 and 0.0056 with a source term, and −0.0081 and −0.0004 for the full adapter, against 0.0148 and 0.0170 for the fixed shrinkage (Fig. 5f). Neither choosing which responses to calibrate on nor measuring more of them changed this: a prospective selection rule added less than 0.003 (Extended Data Fig. 5e), a fixed weight of 0.3 gave the largest gain and a weight of 1 a loss (Extended Data Fig. 5f), and the gain under an oracle weight rose only from 0.0128 at k = 2 to 0.0179 at k = 40 (Extended Data Fig. 5g). With corrections constructed from all held-out responses, the attainable gain rose from −0.0102 to +0.0063 under the fixed shrinkage and from +0.0089 to +0.0148 under an oracle weight chosen in hindsight, as the total number of control cells increased from 20 to 160 (Fig. 5g). At subsampled depth the mixture released by the shared-control gate was inflated by 0.703 and the mixture released by the split-control gate by 0.086 (Extended Data Fig. 5h), and of four forms of the residual the unshrunk one lowered the score at every budget whereas the other three changed it by at most 0.014 (Extended Data Fig. 5i). In this regime, accuracy is limited by the precision of the control estimate.

Extended Data Figs 7 and 8 report earlier comparisons that were scored under shared controls. A logistic release policy, shown as Logistic policy (ER), was compared with parameter updating of a published model (Extended Data Fig. 7a-c); on external study families it did not reduce the fraction of harmful contexts relative to unconditional adaptation (5 of 96 for both), and its mean PCC was lower by 0.013 (0.371 versus 0.384), failing the prespecified safety criterion (Extended Data Fig. 7d). On the shallow plate, a ridge release policy, shown as Ridge policy (Plate 2), had a positive gain in all 15 cell lines, with median gains of 0.329 to 0.379 (Extended Data Fig. 7e). The median context scores were 0.589 to 0.639, against a mean per-drug ceiling of 0.357 implied by split-half reliability, and at k = 5 the lowest context score exceeded the ceiling of every drug (Extended Data Fig. 7f, g). A score above the reliability of the measurement cannot reflect predictive accuracy. All 225 blinded ridge-policy fits selected weight 1.0 (Extended Data Fig. 6h); in this evaluation the policy therefore coincided with unconditional full-amplitude adaptation and gave the same mean gain (Extended Data Fig. 7h). Distributional alignment, separate comparator suites, external study families and a growth endpoint are shown in Extended Data Fig. 8a, b, Extended Data Fig. 8c, d, Extended Data Fig. 8e, f and Extended Data Fig. 8g, h.

The depth law was reproduced in a second dataset. We used a separate nested control-depth ladder in the chemical screen, spanning 20 to 640 cells per half (32-fold) while holding 25 treated cells per condition fixed; the deep-plate ladder instead used 10 to 160 controls per half and 150 treated cells per condition. The product of depth and inflation stayed within 4.1%, 10.6% and 13.5% of each context’s own constant in its three contexts, compared with 0.8% to 11.4% (median 4.9%) for the individual cell lines of the deep plate, and it was exactly zero for the control-free predictor at every depth (Extended Data Fig. 9a, b). The shared term fell by 76% across the ladder and the independent term changed by 5% (Extended Data Fig. 9c).

### The cross-system part of a fibrosis programme localises to scar in human liver

The principle behind the criterion, that what reproduces across independent estimates is what transports, is not specific to perturbation models. We applied it to a fibrosis programme measured in two systems: primary human hepatic stellate cells and the LX-2 stellate cell line, each treated with TGF-beta^34^. We defined the programme as the 438 genes induced more than twofold by TGF-beta in primary cells (log2 fold change > 1) and, using only the response of the same genes in the cell line and no tissue data, derived two gene sets from it. The cross-system set, CORE, comprises the 94 genes that were also induced more than twofold in the line; the primary-cell-only set, pHSC-specific, comprises the 146 genes that were unchanged in the line (absolute log2 fold change < 0.25). The remaining 198 genes belong to neither set. We also decomposed the response of each of the 438 genes into a shared term, the mean of its log2 fold changes in the two systems, and a residual, half the difference between them. The shared term was the larger for the 283 genes with a positive response in the line (the shared-dominant genes, B-dom in the figures), and the residual for the other 155 (the residual-dominant genes, *Δ*-dom).

We then asked where these gene sets are expressed in 15 spatial transcriptomic sections of paediatric human liver^35^ (19,220 spots), scoring each spot relative to the hepatocyte mean of its section. In the example section BA2_1, the CORE set was elevated in scar by a median of +0.437, and the residual-dominant genes by about half as much on the same spots (+0.224; Fig. 6a). The difference persisted within the highest stellate-cell abundance quintile. The cross-system score correlated with the estimated stellate-cell abundance of a spot (Spearman *ρ_s_* = 0.67), but within the highest abundance quintile scar spots still scored +0.185 higher than hepatocyte spots (2,566 and 1,120 spots; one-sided Mann-Whitney *P* = 1.4 × 10^−181^), compared with +0.055 for the residual-dominant genes (Fig. 6b).

**Figure 6.**
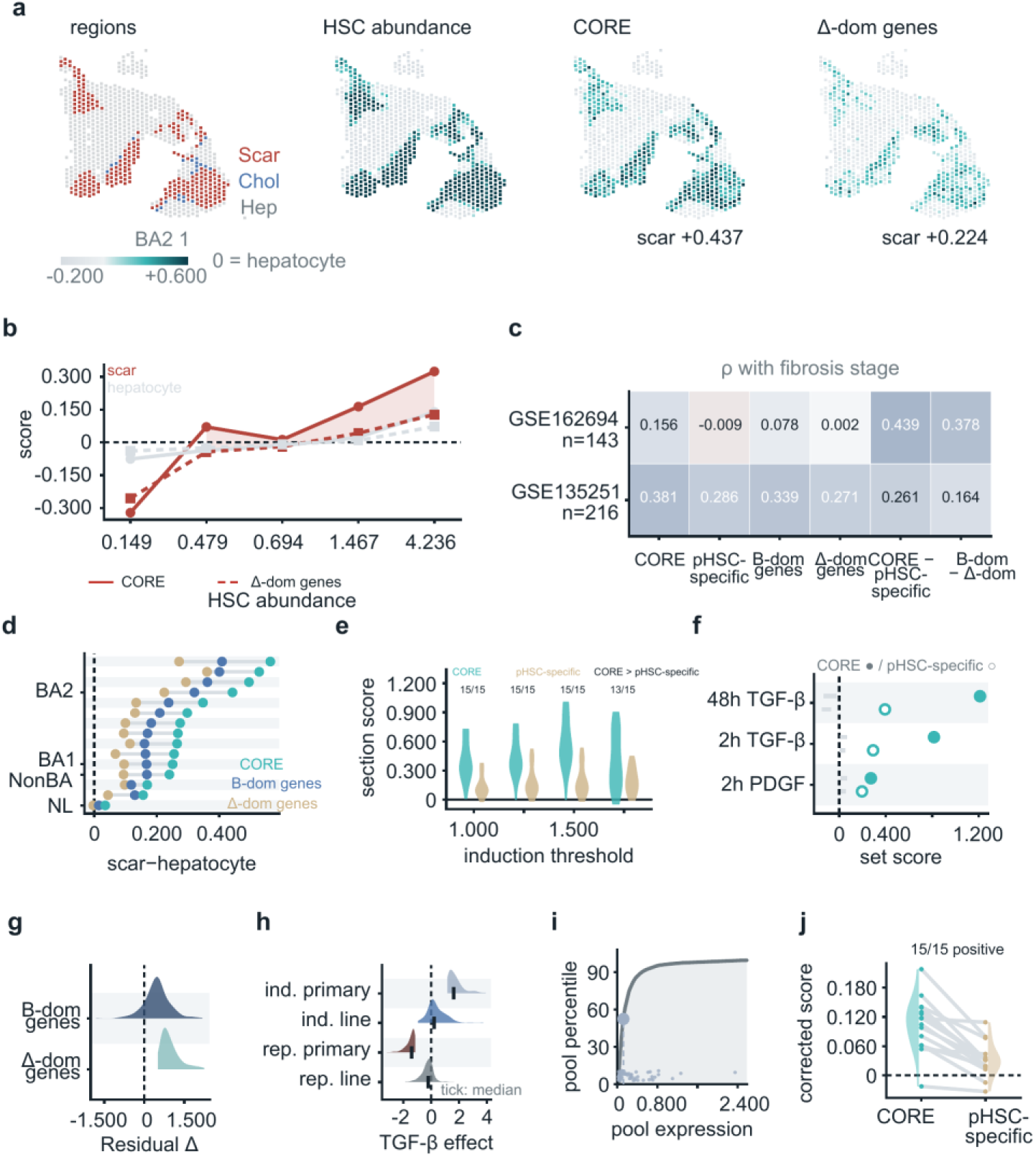
The cross-system part of a TGF-β programme localises to scar in human liver. Gene sets were defined in vitro only. From the 438 genes induced more than twofold by TGF-β in primary human hepatic stellate cells^34^, the cross-system set (CORE; 94 genes also induced more than twofold in LX-2 cells) and the primary-cell-only set (pHSC-specific; 146 genes unchanged in LX-2) were derived. Spatial data are 15 Visium sections of paediatric liver^35^ (19,220 spots); spot scores are referenced to the hepatocyte mean of their section. **a**, One biliary-atresia section: region annotation, stellate-cell abundance from cell2location^40^, and scores for the CORE set and the Δ-dom genes defined in **g**. **b**, Median score by quintile of stellate-cell abundance for scar (red) and hepatocyte (grey) spots; solid, CORE set; dashed, Δ-dom genes. Top quintile, 2,566 scar and 1,120 hepatocyte spots. **c**, Spearman correlation with fibrosis stage in two patient cohorts^36,37^ (*n* = 143 and 216 patients) for the CORE and pHSC-specific sets, the B-dom and *Δ*-dom subsets, and the within-patient differences CORE - pHSC-specific and B-dom - *Δ*-dom. **d**, Scar-minus-hepatocyte contrast per section for the CORE set and the two subsets, with sections grouped on the axis as BA2, BA1, NonBA and NL; BA, biliary atresia; NonBA, non-biliary-atresia liver disease; NL, normal liver. **e**, Section scores of the CORE and pHSC-specific sets at four induction thresholds; counts, sections in which CORE - pHSC-specific is positive (*n* = 15 sections). **f**, Set scores in an independent stellate-cell dataset^39^ under three stimulation conditions against an expression- and size-matched null (500 draws); filled, CORE; open, pHSC-specific; grey bars, null mean to 97.5th percentile. **g**, Residual *Δ* distributions within the B-dom (283 genes) and *Δ*-dom (155 genes) subsets. **h**, log₂ fold change in primary cells and in LX-2 for the 188 induced and 312 repressed genes of the 500-gene activation axis; tick, median. **i**, Expression of the 94 CORE genes against the cumulative distribution of the tissue expressed pool. **j**, Scar-minus-hepatocyte contrast per section after adjustment for cell composition, CORE against pHSC-specific, lines connecting sections.

The association held in two independent patient cohorts^36,37^. The within-patient difference between the cross-system and primary-cell-only sets increased with fibrosis stage in both (Spearman *ρ_s_* = 0.44, *P* = 4 × 10^−8^, n = 143; *ρ_s_* = 0.26, *P* = 1 × 10^−4^, n = 216). Taken alone, neither set was associated with stage in the smaller cohort (0.16, P = 0.06, and −0.01), whereas both were in the larger one (0.38 and 0.29; Fig. 6c). In individual sections, the scar-minus-hepatocyte contrast was positive in all 15 sections for the CORE set and for the shared-dominant genes, and in 14 for the residual-dominant genes (Fig. 6d).

The result did not depend on the threshold that defines the programme. Across four induction thresholds, the within-section difference between the two sets was positive in all 15 sections at the three lower thresholds and in 13 of 15 at the highest (one-sided Wilcoxon signed-rank test^38^, *P* = 3.1 × 10^−5^ and *P* = 0.001; Fig. 6e). In an independent stellate-cell dataset^39^, all six observed values (three stimulus conditions, two gene sets each) exceeded the 97.5th percentile of an expression-matched null (P = 0.002 each, the smallest value attainable with 500 null draws; Fig. 6f).

The division between shared and residual terms is one of degree. The residual was positive for 92% of the 438 genes (Fig. 6g), and both directions of regulation carried over to the cell line at about one seventh of the magnitude: across 500 TGF-beta-responsive genes, the 188 induced genes had a median effect of +1.61 in primary cells and +0.216 in the line, and the 312 repressed genes −1.41 and −0.222 (Fig. 6h). The cross-system genes were not simply the most highly expressed: 68 of the 94 were in the expressed pool of the tissue, with a median expression at the 52nd percentile of that pool (Fig. 6i). After each spot was adjusted for its inferred cell composition using cell2location estimates^35,40^, the difference between the two sets remained positive in all 15 sections (median +0.077), and the CORE set alone in 14 (Fig. 6j). The part of the programme that reproduces across systems in vitro is the part found at the site of disease in tissue.

## Discussion

The standard evaluation of perturbation predictors is inflated by a shared control. Because every response in a context is referenced to one control estimate, the error of that estimate enters both the prediction and the target, and we have shown that the resulting inflation of the unnormalised score is inversely proportional to control depth, and that the inflation of the correlation depends in addition on the control’s noise-to-signal ratio. The inflation was large enough to change the ranking of published methods and to make few-shot adaptation appear beneficial where it was harmful. Splitting the controls removes it and yields a criterion that decides, from calibration data alone, whether a correction should be applied; across six datasets, mean gated gain was non-negative under the original two-control evaluation.

Existing perturbation predictors use latent-space arithmetic, conditional generative models or optimal transport to transfer responses across contexts^8–13^. Their performance is often assessed against the control estimate of the target context. Nicol and colleagues^20^ analysed this form of evaluation. They showed that a control shared between prediction and target inflates correlation-based scores and can change the ranking of methods, that scoring against an independent half of the controls removes the inflation, that the bias depends on control depth, and that the split-control estimator is conservative, attenuated towards zero. Their analysis considers a prediction that is fixed with respect to the target controls. In few-shot adaptation the correction is itself built from responses referenced to those controls, so the control error enters the prediction as well as the score. For this case, we found that the amount of control error a prediction carries varies with the method and its pipeline, from zero for predictors that never read the target controls to more than one when an additive step is attached to a published model. We tested the dependence of the inflation on control depth in nested control-depth experiments, and we used leave-one-out split-control evidence to decide, in advance, whether a correction should be applied in a given context. Systema^16^ addresses a different quantity, the systematic transcriptional variation shared across perturbations in a dataset, rather than the sampling error of a limited control estimate. Related sources of misleading inference include batch effects and unwanted technical variation in bulk expression studies^41,42^, pseudoreplication in single-cell differential expression^43–45^, and data leakage and optimistic evaluation in machine learning^21,22^. Benchmarks of integration and abundance methods likewise show the importance of evaluation design^46–48^.

Two practices follow from these findings. First, a comparison should include at least one method that never reads the target context’s controls. Such a method cannot be inflated and serves as a negative control^49^ that fixes the scale against which the other methods are read^14^; methods that do read the target controls should be expected to lose score when the controls are made independent, and their order should be treated as unresolved until it is recomputed. Second, splitting a context’s controls and scoring against the held-out half requires no new experiment. The difference between the split-control and shared-control scores measures how much of a dataset’s ranking comes from control noise. The same law can be used before an experiment is run: a study can project the control depth associated with a specified bias target under the constant-denominator approximation, and use that projection to plan control allocation.

The principle behind the criterion applies wherever a contrast is defined against a shared reference. In spatial transcriptomics the reference is shared by the spots of a section, and methods that read programmes jointly across cells and spots^50^ or couple expression to tissue morphology^51^ could be tested in the same way. In human liver, the part of a TGF-β programme that reproduced across two in vitro systems was the part that localised to scar, which suggests that reproducibility across independent systems can guide the selection of gene programmes as the cell biology of fibrosis is mapped in more detail^26,52^.

Our study has limitations. The criterion guards against harmful adaptation and is not designed to increase accuracy where adaptation already helps. Its performance relative to other ways of moderating a correction depends on how many control cells a context provides. The criterion identifies whether adaptation helps in a context only after calibration responses have been measured; it does not predict this before any measurement, and it does not rank individual perturbations within a context^15,16^.

In conclusion, this work provides a way to decide, before any held-out response is seen, whether few-shot adaptation should be applied in a new cellular context. The criterion is computed from calibration data alone, leaves the shared predictor unchanged when it abstains and, in the original evaluations across six datasets, avoided the mean losses that unconditional adaptation incurred. It rests on a quantitative account of how a shared control inflates evaluation: a carriage coefficient that describes how much of the control error each method carries, and a law, examined in nested control-depth experiments, that relates the inflation to control depth and the control’s noise-to-signal ratio. As few-shot adaptation becomes a common route for carrying perturbation predictors into new contexts, we expect these tools to make that step safer and the comparison of methods more reliable.

## Methods

### Data

We evaluated the criterion on six datasets from five sources. Tahoe plates 2 and 7^29^ profiled transcriptional responses to drug treatments across human cancer cell lines. sciPlex3^30^ measured responses to small-molecule treatments in human cancer cell lines using multiplexed single-cell transcriptomics. The NeurIPS cross-cell benchmark^15,33^ profiled small-molecule responses across cell types in human peripheral blood mononuclear cells.

McFarland^32^ measured drug responses across pooled human cancer cell lines using MIX-Seq. Haber^31^ profiled mouse small-intestinal epithelial cells under homeostatic and infection conditions. Responses on the shallow plate were computed over 7,070 genes, those shared by three genetic screens (K562, KOLF2.1J and RPE1) and present in the atlas; 7,007 of these genes were matched on the deep plate. The four external datasets were analysed on the 5,000 highly variable genes of their benchmark distribution.

The two Tahoe plates provided the shallow- and deep-control evaluations, and the four external datasets were used to evaluate the criterion across cellular systems. All analyses used publicly available data. No participants were recruited and no new human or animal samples were collected for this study.

### Responses, scores and the two control regimes

A context was the unit to which a prediction was transported: a cell line, a patient sample or a tissue. For Tahoe, counts were normalised to 10,000 per cell and log1p-transformed; external expression matrices were analysed on the scales supplied by their respective pipelines. Cell profiles were averaged within each condition to form a pseudobulk profile. A perturbation response was the treated profile minus the control profile. For each perturbation within a context, we calculated the Pearson correlation between predicted and observed responses across genes. Per-condition aggregation avoids treating individual cells as independent replicates^43–45^.

Supplementary Table 1 records the control definitions and splitting levels for the criterion evaluation, the benchmark re-analysis and the candidate archives. Atlas controls were vehicle cells from the same cell line and plate, split at cell level. The chemical datasets used vehicle wells or cells, split by plate for sciPlex3, by plate_name for the NeurIPS criterion evaluation and by condition3 for McFarland. Haber used uninfected controls, split by batch. The candidate archives used library_id for NeurIPS and channel for McFarland. Using these dataset-specific keys, we divided the control cells into two disjoint halves with equal numbers of cells, denoted A and B. Predictions and calibration responses were constructed with half A. Shared-control scores used responses referenced to A; independent-control scores used responses referenced to B, with predictions, treated cells and calibration sets held fixed.

Analyses with two folds rebuilt and scored each exchanged A/B direction separately and then averaged the two values; the atlas-plate criterion used the single A-to-B direction specified below.

The theoretical noise-to-signal ratio was defined as

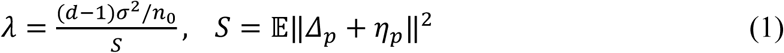

where *d* is the number of genes, *n*_0_ the number of cells in one control estimate, *σ*^2^ the per-gene control variance, and *Δ_p_* + *η_p_* the response including treated-cell sampling noise. The model-free screening ratio reported in Fig. 4a-d differs from equation (1) in its denominator: its numerator was half the centred squared distance between the control half-means, and its denominator averaged the inner product of the two responses to each perturbation, each referenced to a different control half, with the deposited screening implementation using uncentred response vectors. Estimates were summarised by the median over contexts and splits, and estimates from the calibration perturbations alone are compared with estimates from all perturbations of a context in Extended Data Fig. 1e, f. The dataset-wide survey used all eligible contexts and perturbations in each deposit; benchmark-specific analyses used the contexts and perturbations scored by the corresponding pipeline.

The carriage coefficient was estimated by constructing a prediction from each control half and projecting their difference, *ΔP*, onto *Δc* = *c_A_* - *c_B_*: *a* = −⟨*ΔP*, *Δc*⟩/‖*Δc*‖^2^. Both vectors were centred over genes, and the cosine between them was also reported. The norm ratio ‖*ΔP*‖/ ‖*Δc*‖ equals the directional coefficient when *ΔP* = −*aΔc* with *a* ≥ 0. The twelve benchmark estimates compared calibration and target predictions; the two Tahoe estimates used full target-response records. A predictor independent of target controls has *a* = 0. Adding the mean calibration response at unit weight to such a predictor gives *a* = 1. For components using the same target-control estimate, a sum with fixed weights has coefficient *a* = ∑*_j_w_j_a_j_*, where *w_j_* and *a_j_* are the weight and coefficient of component *j*.

### Shared-control inflation

Let *T_p_* denote the measured profile under perturbation *p* and *μ*_0_ the mean profile of *n*_0_ control cells. The observed response was written as

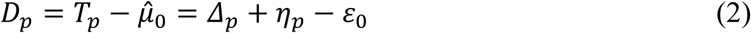

where *Δ_p_* is the true response, *η_p_* the treated-cell sampling noise and *ε*_0_ the error of the shared control estimate. Under the scalar-carriage model, the prediction is

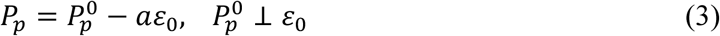

with a control-independent component *P_p_*^0^ and a fixed carriage coefficient *a*. Under isotropic control noise with per-gene variance *σ*^2^, Proposition 1 compares the expected centred inner products obtained with shared and independent controls:

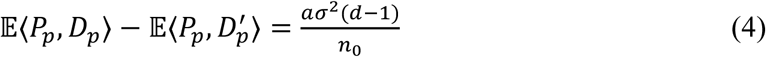

Here *D_p_*’ uses an independent control estimate. Gene centring gives E‖*ε*_0_‖^2^ = (*d* - 1)*σ*^2^/*n*_0_. When the centred prediction difference is *ΔP* = −*aΔc*, exchanging the control halves gives the two-fold identity of Proposition 1’,

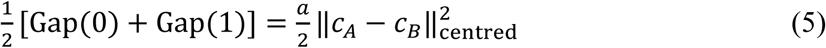

where *c_A_* and *c_B_* are the control half-means. Here Gap(0) = ⟨*P_A_*, *D_A_*⟩ - ⟨*P_A_*, *D_B_*⟩ and Gap(1) = ⟨*P_B_*, *D_B_*⟩ - ⟨*P_B_*, *D_A_*⟩, using the same treated profiles. With the projection estimate *a* in place of *a*, equation (5) is a sample identity for the unnormalised inner-product gap rather than an independent test of the model. On the correlation scale, Proposition 2 gives

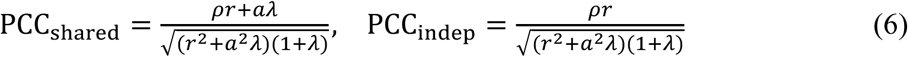

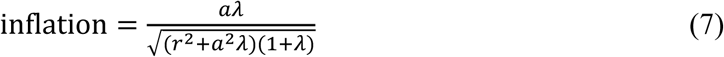

where *ρ* is the correlation of the control-free prediction with the response, *r* their scale ratio and *λ* the ratio in equation (1). These expressions describe ratios of second moments, with matched control-error variance and with control errors independent of the control-free prediction and of the response; equations (6) and (7) were evaluated per perturbation before averaging, and mean realised PCC was averaged separately. For a linear control map *M* with control-error covariance *Σ*, the expected inner-product gap is tr(*MΣ*), with *M* and *Σ* acting on gene-centred vectors; the scalar model takes *M* = *aI*. In simulations covering 12 settings over a 52-fold range of *λ*, we compared equation (7) with the second-moment ratio and with inflation calculated from mean per-perturbation correlations. Supplementary Note 1 gives the derivations of Propositions 1, 1’ and 2 and of the control-depth condition, together with the design and results of these simulations.

For the signal-normalised inner-product gap, the control-depth condition was

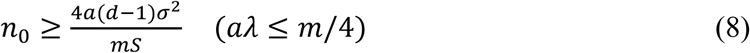

where *m* denotes a reference margin on the signal-normalised inner-product scale, with the allowed bias set to one quarter of that margin. In the separate PCC-based benchmark projections, the reference margin was the observed gap between adjacent methods, with bias targets of one quarter or one tenth of that gap. For PCC-based depth estimates, we approximated the denominator in equation (7) as constant, so that inflation scaled inversely with control depth, and the projections in Fig. 2g used the observed PCC inflation rather than equation (8).

### The criterion

The shared predictor is the mean response to the same perturbation in the other contexts: for a target context *c* and a perturbation *p*, 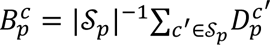, where S*_p_* is the set of contexts other than *c* in which *p* was measured and each *D_p_*^c’^ is formed against the control of its own context. It has no fitted parameter and reads no cell of the target context, so its carriage coefficient is zero; figures and tables call it Pooled. *B_q_* denotes this prediction, whereas the superscript B in *D_q_*^B^ denotes control half B. An adapted prediction is

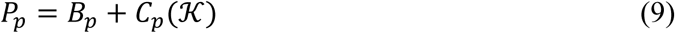

where K is the calibration set, the *k* perturbations of the target context whose responses were measured, and *C_p_* the correction an adapter derives from those responses. Abstaining sets *C_p_* = 0.

We used two adapters. The offset adapter adds the mean calibration residual under a weight *w*, 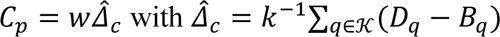, and was applied with *w* = 1 on the external datasets. The mixture adapter, used on the atlas plates, combines four candidate predictions, 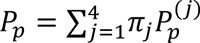: the shared predictor, 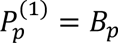; the full offset, 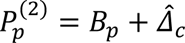; a weighted combination of the responses of the other contexts, 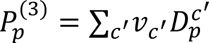; and a low-rank offset, 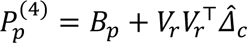, in which the columns of *V_r_* are the first *r* right singular vectors of the other contexts’ responses after the mean over those contexts is subtracted for each perturbation, and *r* is the rank among 4, 8 and 16 with the lowest squared error on the calibration responses. The weights *π* and *v* are non-negative and sum to one; each set was fitted to the calibration responses by least squares with a ridge penalty of 0.01 towards equal weights.

The criterion is the gain of the adapter on the calibration responses when the control that builds the correction and the control that references the scored response are different halves of the target context’s control cells:

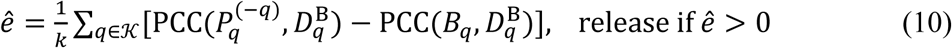

where PCC is the Pearson correlation over genes, *P_p_^(-q)^* the prediction for calibration perturbation *q* of the adapter fitted to the other *k* - 1 calibration responses, all referenced to control half A, and *D_q_*^B^ the observed response of *q* referenced to control half B. The two halves share no cell, and no held-out response is read. The shared-control evidence is the same quantity with *D_q_*^A^ in place of *D_q_*^B^. On the atlas plates we computed the criterion with the adapter built on half A and scored against half B; on the external datasets we also rebuilt and scored each exchanged direction and averaged the criterion values and the held-out gains. The candidate that is released is the complete adapter fitted to all *k* calibration responses referenced to half A. The correction is released when *e* > 0 and withheld otherwise, without a tuned parameter. In the original evaluation, the held-out gain of a trial is the mean, over the perturbations outside the calibration set, of the difference in correlation between the adapter fitted to all *k* calibration responses and the shared predictor, both scored against responses referenced to control half B.

We compared the zero threshold with thresholds selected by leave-one-context-out validation on the deep plate (Fitted-threshold gate): each threshold was selected on the remaining contexts and evaluated on the held-out context. We also evaluated transfer of the selected threshold to the shallow plate. Weight comparisons used the deep plate at native depth, including the 15 cell lines of the criterion evaluation and SNU-423, and the 86 drugs with at least 400 cells in every line. We drew 20 calibration sets per cell line at each budget (*k* = 5 and *k* = 20). We compared an offset weight fitted by leave-one-out on the calibration responses, a simplex-weighted combination of four candidate corrections regularised towards equal weights at three regularisation strengths, and a fixed weight of 0.3. Performance was evaluated on held-out responses; an oracle weight selected on those responses provided a reference.

The criterion, the two adapters and the estimators of the carriage coefficient and the noise-to-signal ratio are implemented in SafeDelta, an open-source Python package that depends only on NumPy. Given the responses of the other contexts, the calibration profiles of the target context and the mean profiles of its two control halves, SafeDelta returns the criterion, the release decision and the predictions for the remaining perturbations.

### Experiments on the atlas plates and external datasets

The Tahoe criterion evaluation used 15 cell lines and 88 drugs per plate. Plate 2 was selected from metadata before expression values were read. Plate 7 used its 15 most abundant lines (SW480, NCI-H460, PANC-1, Hs 766T, MIA PaCa-2, HOP62, SHP-77, A498, CFPAC-1, LoVo, LOX-IMVI, J82, A549, NCI-H2347 and RKO) and drugs at 0.05 µM with at least 400 cells per line. Plate 2 used 20 treated cells per condition and ten vehicle cells per control half; plate 7 used 400 treated cells and 400 vehicle cells per half.

On each Tahoe line, we fitted the mixture adapter using five calibration drugs and evaluated the remaining 83, with eight calibration draws on plate 2 and ten on plate 7. Separate blinded runs, reported as Ridge policy (Plate 2), used 15 contexts, three budgets and five seeds, with fits fixed before held-out outcomes were read (Extended Data Fig. 6h). On sciPlex3, NeurIPS, McFarland and Haber, we used the offset adapter at *w* = 1 and three random partitions of the dataset-specific control keys. Each external scored response was defined by a context, a held-out perturbation and a partition. Calibration used five responses when at least six perturbations were available; otherwise it used one fewer than the available perturbations, with a minimum of two.

For the additional-control evaluation on the deep plate, predictions, release decisions, calibration and test perturbations, and treated-cell profiles were held fixed. Three additional groups, C0-C2, each contained 400 control cells and were disjoint from the original A and B groups and from one another. They were used only for final scoring after predictions and release decisions were fixed (Extended Data Fig. 2a-d). A and B each contained 400 controls, 800 together; adding one C group brings the context total to 1,200 and all three to 2,000, shared across the drugs of a context. Separate fixed-total audits divided the same budget three ways, at 20/20/20, 80/80/80 or 268/266/266 cells. External audits used cell-disjoint or key-disjoint splits with three control draws and ten calibration draws, and held-out gains were averaged within context and draw. The strategy comparisons covered split leave-one-out, its equivalent direct split rule, shared leave-one-out, always adapting, never adapting, a fixed half amplitude, and leave-one-out selection among amplitudes 0, 0.5 and 1.

Adapter comparisons used the same 15 plate 7 lines at native depth and after subsampling to the first 20 treated cells and first ten cells of each control half, with twelve calibration draws per line. Comparators were the shared predictor, scalar rescaling with or without an intercept, full offset, rank-16 offset, and mixture adaptation released by the shared-control or the split-control gate. Budget comparisons and deep-plate model runs used a separate sixteen-line, thirty-drug subset: budgets of two, three and five responses were evaluated on six lines (five at *k* = 3), with up to 500 vehicle cells divided equally, three control partitions and all non-calibration drugs held out.

The control-depth sweep used the 15 plate 7 lines plus KATO III and thirty drugs. Control halves were nested-subsampled to 10, 20, 40, 80 and 160 cells, holding the same 150 treated cells per condition fixed. Each line used five calibration and 25 held-out drugs, with both directions of control splitting; the shared predictor averaged the other fifteen lines at each depth. We evaluated the shared predictor, the calibration mean, and the shared predictor plus the mean calibration response at unit weight. A separate weight-and-capacity analysis used noncumulative control subsets totalling 20, 40, 80 or 160 cells across the two halves; corrections were constructed from five calibration responses or, for the attainable ceiling, from all held-out responses. Under the oracle weight, the gain of the five-response correction rose from 0.0080 to 0.0103 and the ceiling from 0.0089 to 0.0148 across these depths, whereas a correction built and scored with the same evaluation controls gained 0.84 at 20 cells and 0.46 at 160 (Extended Data Fig. 1g). The score of the frozen predictor rose from 0.129 to 0.383 over the same range (Extended Data Fig. 1h).

The sciPlex3 depth replication used disjoint groups of whole plates, 20, 40, 80, 160, 320 or 640 cells per control half, 25 fixed treated cells and the same three estimators. We assessed variation across three control draws at fixed depth and used sign tests to assess changes in depth-scaled control-error energy between the lowest and highest depths.

Prediction accuracy was measured by Pearson correlation across genes on held-out responses. We evaluated the release rule by its gain over the shared predictor, assigning zero gain to withheld trials, the fraction of trials released and the fraction of released trials with positive gain. Depth analyses compared shared- and independent-control scores and assessed the inner-product gap multiplied by control depth against pooled and context-specific constants:

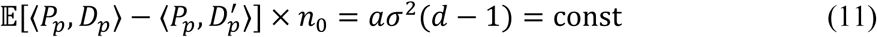

### Published models and benchmark re-analysis

We ran trVAE^10^, scGen^9^ and scPRAM^12^ using benchmark settings and log-normalised expression of 5,000 highly variable genes. trVAE used scArches, two 128-unit hidden layers, a ten-dimensional latent space, dropout 0.2 and mean-squared-error loss; training used learning rate 0.001, batch size 512 and early stopping, for up to 1,000 epochs on the four benchmark datasets and 120 epochs in the Tahoe and sciPlex3 runs, where few-shot fine-tuning on source contexts and calibration responses used 40 epochs and learning rate 0.0001. scGen used batch size 64, early-stopping patience 25 and up to 200 epochs on the benchmark datasets, and batch size 256, patience ten and up to 60 epochs on Tahoe; scPRAM 0.0.2 used 100 epochs and ratio 0.005. Models received one control half and were scored against responses referenced to each half.

For adjacent methods in published leaderboards^14–16^, we calculated paired context-perturbation score differences, 95% context-clustered intervals and shared-control inflation, taking the larger of the two individual-score inflations in each pair. Under the constant-denominator approximation, control depth was multiplied by the observed PCC inflation divided by 25% or 10% of the reported margin. We implemented the pooled and calibration-mean baselines; additional leaderboard methods were assessed from reported scores.

We compared five arms on matched trials, using five seeds and calibration budgets of one, two, three and five responses. ER baseline is the frozen prediction of each original package. ER always-full applies the adaptation of that package at full amplitude in every trial. Logistic policy (ER) releases the adaptation at an amplitude chosen by a logistic classifier of harmful outcomes, specified below. The remaining two arms are full fine-tuning and joint retraining. These comparisons used the shared-control scoring of their original runs. Fine-tuning and retraining were restricted to two dataset families with updatable checkpoints. The logistic policy classified harmful outcomes, defined as a PCC gain below −0.01, at amplitudes 0.25, 0.5, 0.75 and 1, using family-balanced weights, median imputation and standardised calibration features, and was trained with the evaluated study family held out. Inner-family validation selected probability thresholds and amplitudes, retaining candidates with coverage of at least 0.2, non-negative mean gain and a harmful-context fraction no higher than always-full. Metrics were changes in Pearson correlation and cosine similarity, correlation over the top hundred differentially expressed genes, energy distance^53^ and sliced Wasserstein error.

The external-family evaluation of the logistic policy used five study families at a calibration budget of two, averaging five seeds per context. A separate ridge policy, Ridge policy (LOFO), predicted gains at four nonzero amplitudes from nine calibration features: the cross-fit gain mean, minimum, standard error and sign agreement, residual-direction agreement, the candidate-to-baseline magnitude ratio, expert disagreement, source-support distance and the calibration count. It used median imputation, standardisation and a family-balanced multi-output ridge, with inner-family validation selecting the regularisation; the largest predicted positive gain selected the amplitude against zero, ties favouring the lower amplitude. The evaluated family was omitted, and permutation nulls shuffled weights globally or within family on the matched gain table. Ridge policy (Plate 2) is the ridge policy applied to the shallow plate, with router-trained models for each budget and fixed prediction packages. In the comparison of release policies on that plate, V4 always-full and V4 always-half apply the correction of the same packages in every context at full and at half amplitude, the train-only weight is a single amplitude chosen on the training contexts, and the oracle weight is chosen on the held-out responses (Extended Data Fig. 7). The MIX-seq comparator suite used 47 contexts, six test targets and two calibration targets on 1,199 genes, with simplex ridge 0.01 and rank-eight fingerprint ridge. Growth validation used a genome-scale K562 screen^54^ with five held-out-target folds and independently measured growth phenotypes excluded from training and selection. Genetic screens used non-targeting guides as controls without control splitting.

### Analysis of hepatic stellate-cell responses and liver fibrosis

In GSE253493^34^, the TGF-beta-treated triplicates were S4-S6 for primary human hepatic stellate cells and S10-S12 for LX-2; controls were S1-S3 and S7-S9. Condition assignments follow the official GEO solvent and TGF-beta labels and the executed sample-group table. Gene sets were defined within the source table of symbol-mapped genes with primary-cell log_2_FC > 1. The cross-system set, CORE, additionally required LX-2 log_2_FC > 1; the primary-cell-only set, pHSC-specific, required LX-2 |log_2_FC| < 0.25.

For each gene, the shared term was the mean log_2_FC across the two systems and the residual half their difference, primary cells minus LX-2. Within the source table, genes were assigned to the shared-dominant set (B-dom) when the LX-2 response was positive and to the residual-dominant set (*Δ*-dom) otherwise. The activation axis used the 500 genes with the largest absolute primary-cell responses and their matched effects in LX-2.

Spatial analysis used 15 paediatric liver sections from GSE338525^35^. Within each section, genes with mean log1p-CP10k > 0.02 were z-scored across spots, and set scores averaged those gene scores. Scores were referenced to the section’s hepatocyte mean; the section statistic was median(scar) minus median(hepatocyte). Composition adjustment used the cell2location estimates^40^ deposited with the spatial dataset^35^. External stellate-cell validation used GSE119606^39^, and patient-cohort analyses used GSE162694^36^ and GSE135251^37^.

Null gene sets were matched by size and decile of mean expression, with 300 draws for tissue threshold sensitivity and 500 for external validation, for which the smallest attainable empirical P value is 1/501. Threshold sensitivity redefined and rescored both sets at induction cutoffs of 1, 1.25, 1.5 and 2.

### Statistical analysis

Paired section- or context-level comparisons used one-sided Wilcoxon signed-rank tests^38^; intervals were obtained by context-clustered bootstrap^55^. Scored responses and spots within a context or section were not independent; spot-level Mann-Whitney results describe within-section separation. Per-condition pseudobulk aggregation avoided cell-level pseudoreplication^43–45^. Other analyses used Spearman correlation, sign tests and expression-matched random-set nulls. Figure 3f used 4,000 hierarchical bootstrap resamples over three contexts and six context-prediction-replicate units, retaining the original row weighting over 471 scored responses per regime; the plotted intervals are 95% percentile intervals.

For the additional-control evaluation, test-perturbation gains were averaged within each trial and then within context, with equal context weights and zero gain assigned to withheld trials. The 95% percentile intervals used 4,000 context-bootstrap resamples. Evidence–gain correlations used ungated candidate gains; repeated C references rescored the same 150 deep-plate decisions.

## Supporting information

Supplementary Table 1: control definitions

## Acknowledgements

This work was supported by the Natural Science Foundation of China (T2600420) and the China Postdoctoral Science Foundation (2026M794302 and GZC20262144).

## Author contributions

W.-L.L., Y.-B.Y. and Q.-Y.Z., together with W.-Y.S., Y.-F.C. and R.-R.H., conceived and designed the study. W.-L.L. and Y.-B.Y. developed the methodology. Y.-B.Y. was responsible for code development and organisation. Q.-Y.Z., J.-C.F., Q.-F.W., Y.L. and S.-Y.W. contributed to data curation and analysis. W.-L.L. and J.-L.T. contributed to computational methodology and validation. J.D., Y.B. and Y.-F.L. contributed to study conceptualisation and critically reviewed the manuscript. W.-L.L., Y.-B.Y. and Q.-Y.Z. prepared the figures and drafted the manuscript. W.-Y.S., Y.-F.C. and R.-R.H. supervised the study and critically revised the manuscript. All authors contributed to writing and revising the manuscript and approved its final submission.

## Data availability

Tahoe-100M²⁹ is available at https://huggingface.co/datasets/tahoebio/Tahoe-100M. Processed matrices for sciPlex3³⁰, McFarland³², Haber³¹ and the NeurIPS cross-cell benchmark³³, in the partition distributed with a published benchmark¹⁵, are available at https://doi.org/10.5281/zenodo.14607156. The five pathway screens⁵⁶ are available at https://doi.org/10.5281/zenodo.14518762 (raw data, GEO accession GSE281048) and the genome-scale K562 screen⁵⁴ at https://doi.org/10.25452/figshare.plus.20127869.v1. The Afriat dataset⁵⁷ and the candidate datasets that were examined but not admitted⁵⁸⁻⁶⁰ are part of the same benchmark archive. The liver datasets are available from GEO under accessions GSE253493³⁴, GSE338525³⁵, GSE119606³⁹, GSE162694³⁶ and GSE135251³⁷.

## Code availability

The source code for SafeDelta and the accompanying analysis scripts are available at https://github.com/yyb2020/safedelta. SafeDelta is also available as a Python package on PyPI at https://pypi.org/project/safedelta/.

---

**Extended Data Fig. 1.**
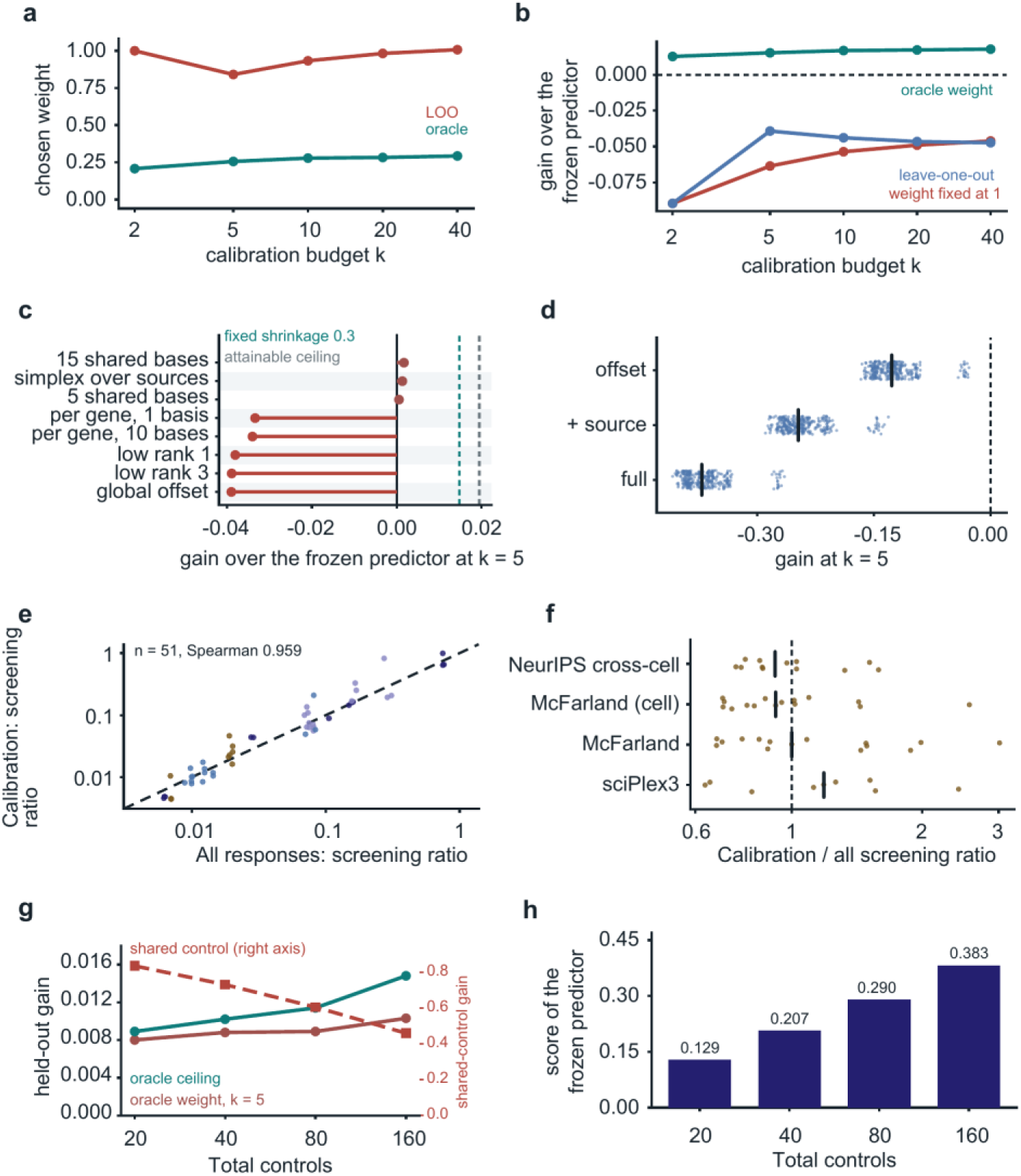
Adaptation weight, adapter capacity and control depth on the deep plate. Deep plate unless stated. **a**, Adaptation weight chosen by leave-one-out (LOO) on the calibration responses and the oracle weight chosen on held-out responses, against calibration budget *k* = 2 to 40. **b**, Held-out gain over the frozen predictor against *k* for three weight rules: oracle, LOO, and a weight fixed at 1. **c**, Held-out gain at *k* = 5 for eight adapter capacities (global offset, low-rank offsets of rank 1 and 3, per-gene offsets with 1 and 10 bases, 5 and 15 shared bases, simplex over sources); dashed lines, fixed shrinkage of 0.3 and the capacity ceiling attainable at this budget. **d**, Held-out gain at *k* = 5 for three adapter rungs with the weight fixed at 1, one point per context and calibration draw (16 cell lines, 20 draws each; *n* = 320); bar, median. **e**, Screening ratio computed from the calibration perturbations alone against the ratio computed from every perturbation of the context, with the same control cells in both (*n* = 51 context-by-split estimates; Spearman 0.959); dashed line, identity. **f**, Ratio of the calibration-only to the all-perturbation screening ratio per dataset, with McFarland shown under both of its split keys; bar, median; dashed line, ratio of 1. **g**, Held-out gain under the oracle weight against total control cells (20–160), for a correction built from *k* = 5 calibration responses and the ceiling from a correction built from all held-out responses (left axis); right axis, gain from a correction built and scored using the same evaluation controls. **h**, Absolute PCC of the frozen predictor at each control depth.

**Extended Data Fig. 2.**
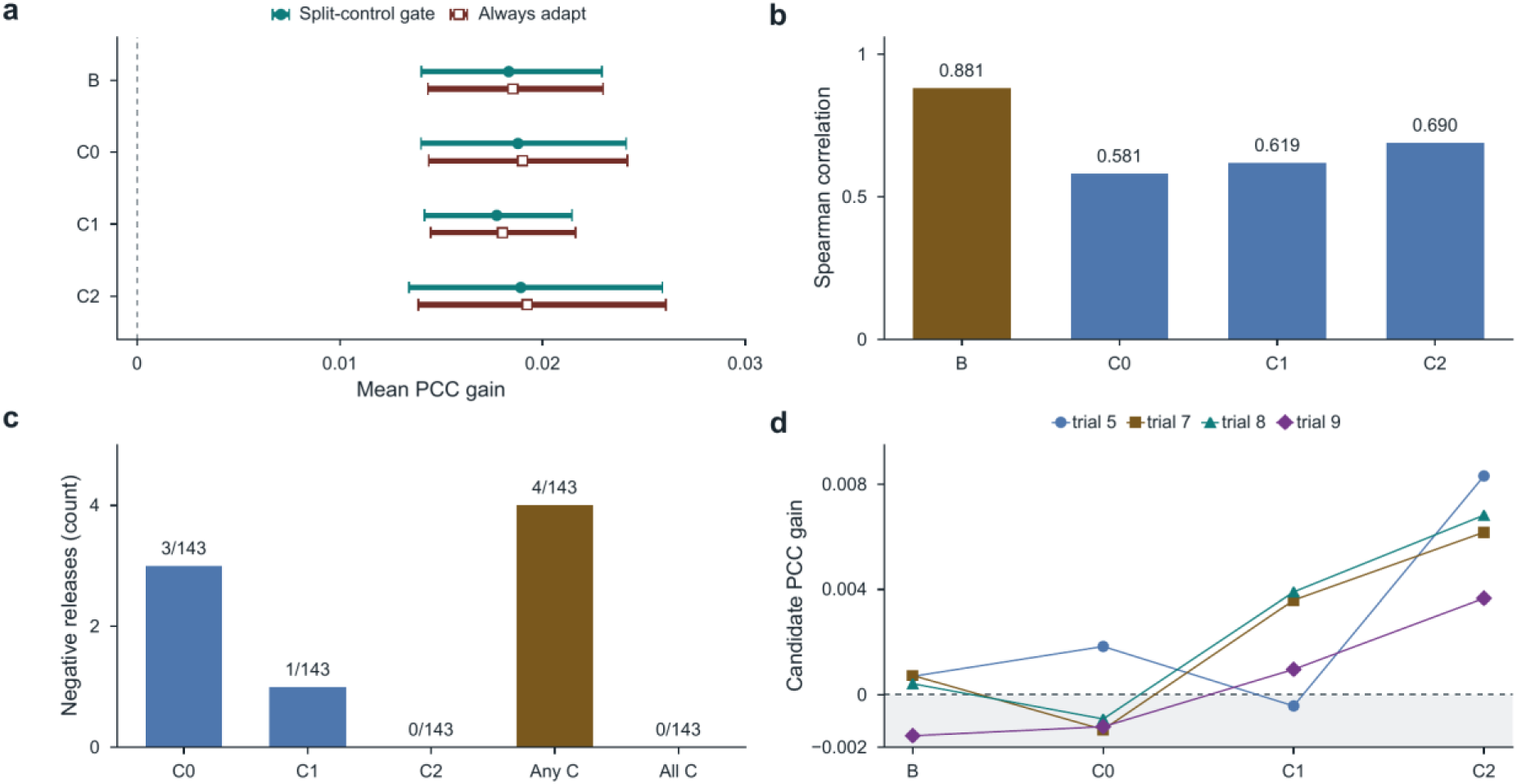
Independent-control evaluation of fixed adaptation decisions on the deep plate. **a**, Mean PCC gains under gated adaptation (filled circles) and unconditional adaptation (open squares). B is the original decision reference; C0–C2 are additional, mutually disjoint evaluation references. Bars, 95% context-bootstrap intervals. **b**, Spearman correlations between the original criterion and ungated candidate gains, using the same 150 trials under each reference. **c**, Released trials with negative mean test gains under each C, any C or all C; the denominator is the same 143 releases. **d**, Paired gains across references for all four sign-unstable releases, from CVCL_0023. The evaluation used 15 contexts with ten calibration draws each. Predictions and release decisions were fixed before scoring against C; A, B and each C group contained 400 cells and shared none. Means include all 150 trials, with zero gain assigned to withheld trials.

**Extended Data Fig. 3.**
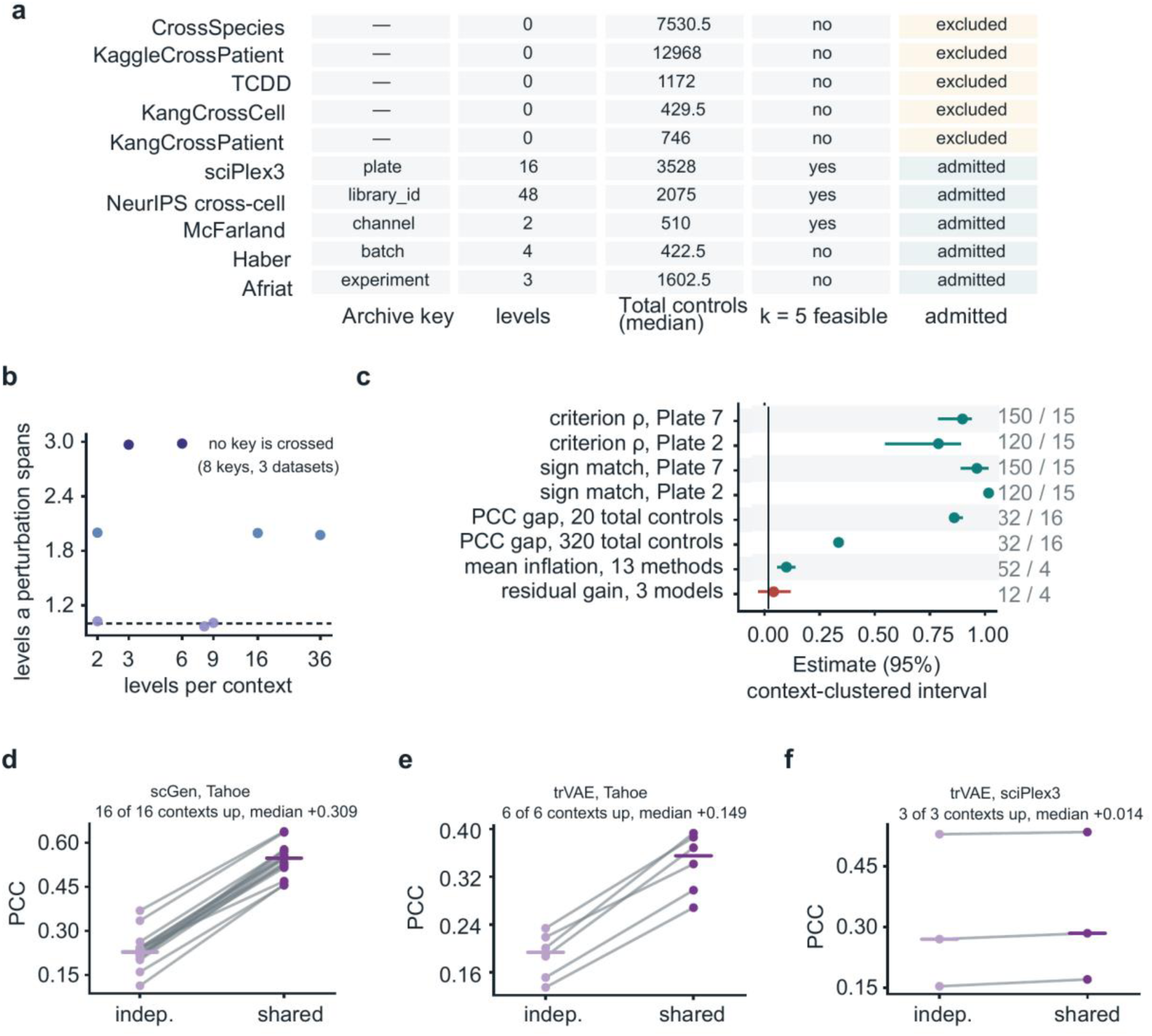
Dataset admissibility, context-clustered intervals and published models under the two control regimes. **a**, Candidate-dataset audit: archive split key, number of key levels, median total control cells, feasibility of *k* = 5 and inclusion status for ten candidates^58–60^. **b**, Number of key levels per context against the number of levels a perturbation spans, for the eight candidate split keys across three datasets; dashed line, one level per perturbation. **c**, Point estimate and 95% percentile interval from 2,000 resamples of whole contexts or datasets, according to each row’s clustering unit^55^, for eight claim-by-stratum rows. The number of observations and clusters is given beside each row. **d-f**, Per-context mean PCC under independent and shared controls, one line per context: scGen on Tahoe (*n* = 16 contexts), trVAE on Tahoe (*n* = 6) and trVAE on sciPlex3 (*n* = 3); bar, median.

**Extended Data Fig. 4.**
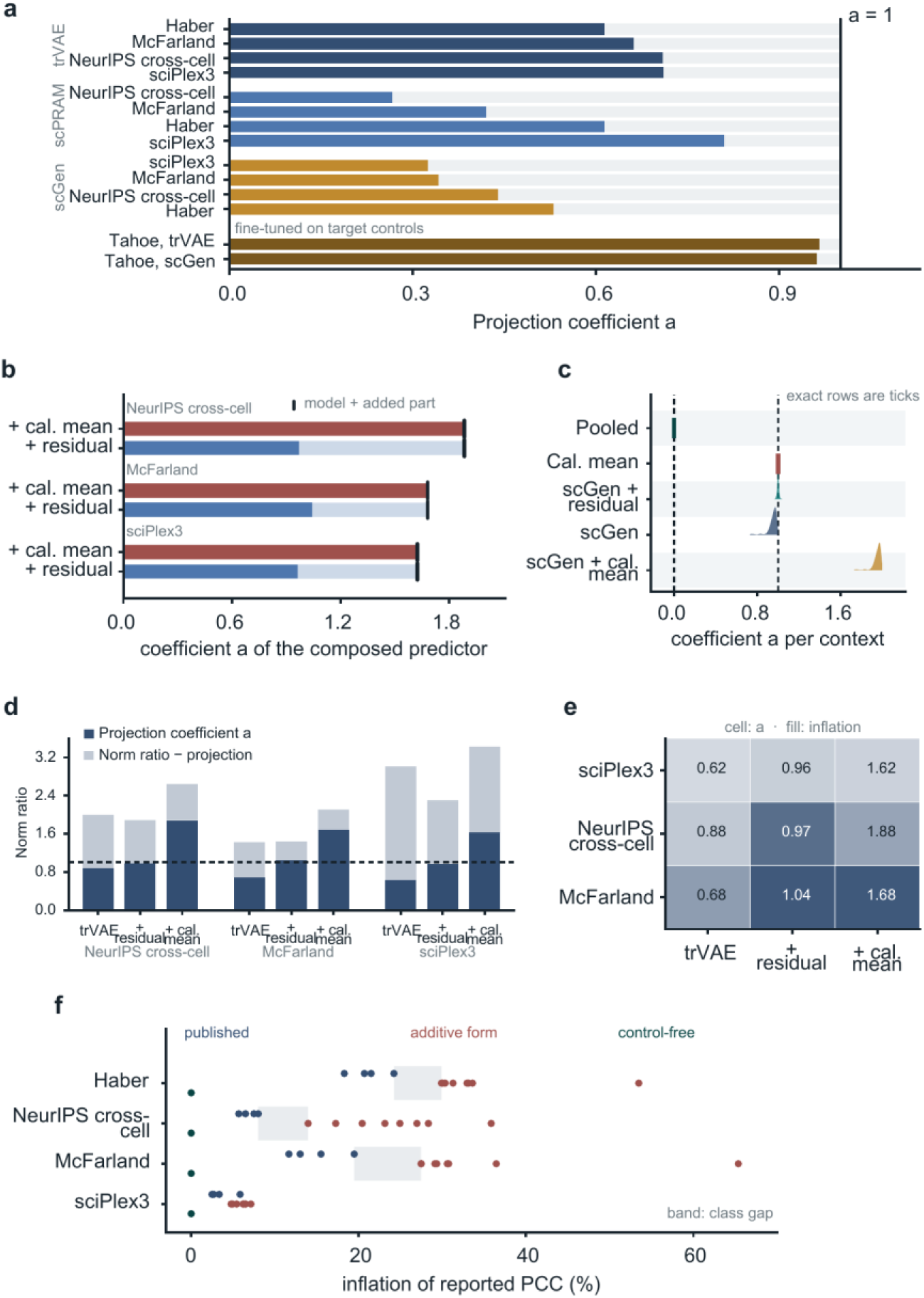
Estimation of the carriage coefficient. **a**, Projection coefficient *a* of each published predictor, averaged over the calibration perturbations, four datasets per model, with the two models fine-tuned on target-context controls shown separately; dashed line, *a* = 1, the value of a unit-weight additive calibration mean. **b**, Coefficient of a composed predictor (published model plus calibration mean, or plus offset) against the sum of its components’ coefficients (tick) for three datasets. **c**, Per-context distribution of *a* at native depth for five predictors (*n* = 1,050 context-target pairs per predictor). **d**, Projection estimate of *a* (dark) and the excess of Norm ratio over it (light), with total bar height equal to Norm ratio, for three predictors on three datasets; dashed line, *a* = 1. **e**, Coefficient *a* (printed) and inflation of the reported PCC (fill) for the same predictors and datasets. **f**, Inflation of the reported PCC for the three predictor classes (published, additive form, control-free) on four datasets; band, gap between classes.

**Extended Data Fig. 5.**
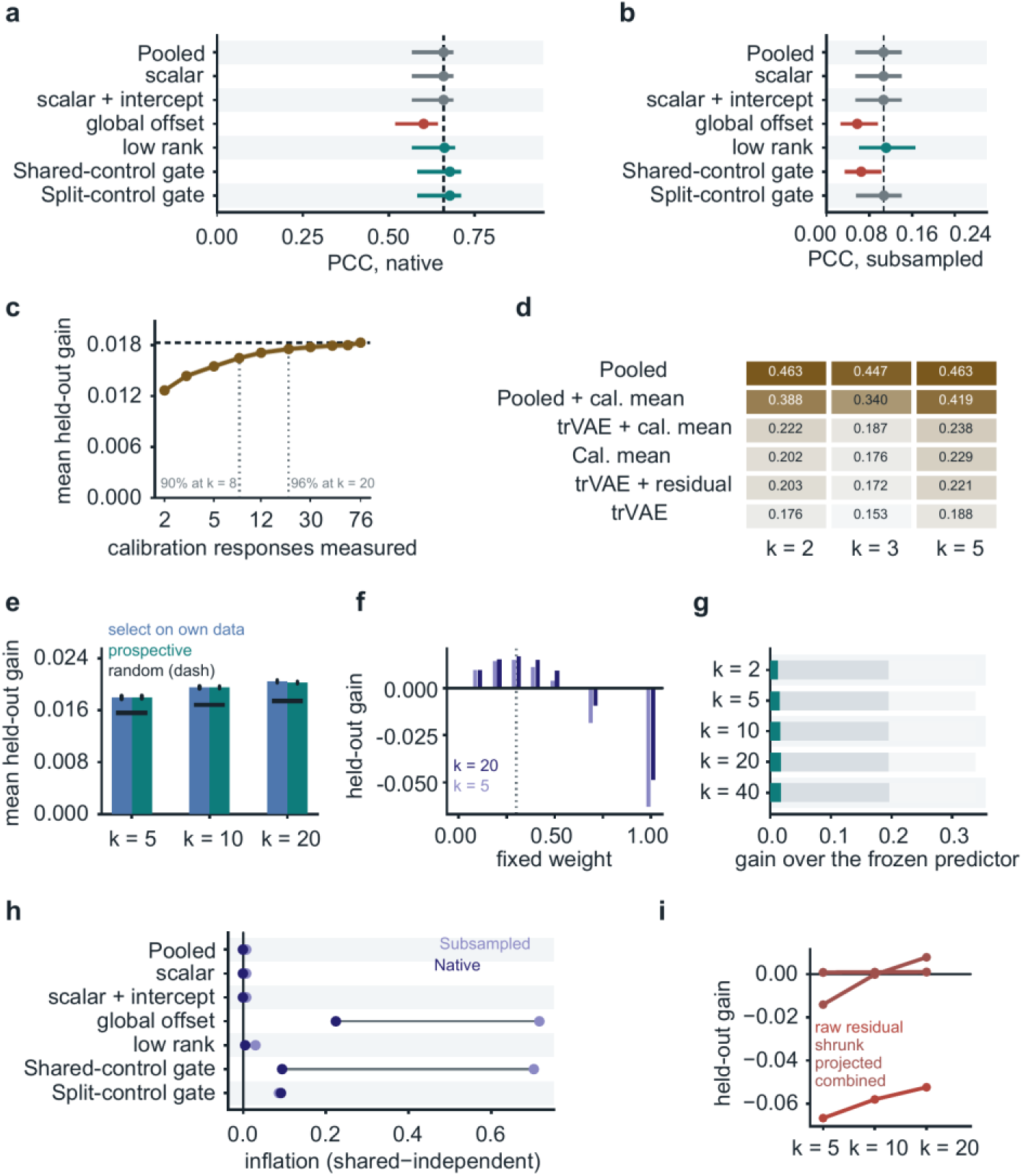
Calibration budget, response selection and adapter comparison. Deep plate throughout. **a-b**, Independent-control PCC of each rung at native depth (a; 400 treated cells per condition and 400 controls per half) and subsampled depth (b; 20 treated cells and ten controls per half); point, mean over 180 trials; bar, range; dashed line, Pooled. **c**, Mean held-out gain against calibration budget, 2 to 76 responses; ribbon, ± 1 s.e.m.; dotted lines, budgets at which 90% and 96% of the gain at 76 responses is reached. **d**, Independent-control PCC of six methods at *k* = 2, 3 and 5 on six cell lines (five at *k* = 3), controls split into halves of 250 cells (198 and 199 in one line); the run behind Fig. 2b,f and Fig. 3h. **e**, Mean held-out gain at *k* = 5, 10 and 20 when calibration responses are chosen at random (dashed), by selection on the context’s own data, or by a prospective rule; bars, 95% interval of the difference from random. **f**, Held-out gain for fixed weights 0 to 1 at *k* = 5 and *k* = 20; dotted line, weight 0.3. **g**, Oracle-weight gain of a correction built from calibration responses at *k* = 2 to 40, stacked with the additional gain from a context-oracle weight and a full-oracle correction; all oracle choices use held-out responses. **h**, Mean inflation (shared minus independent PCC) of each rung over the 180 trials at native and subsampled depths; Shared-control and Split-control gates are separate rungs. **i**, Held-out gain of four residual forms (raw, shrunk, projected, combined) at *k* = 5, 10 and 20.

**Extended Data Fig. 6.**
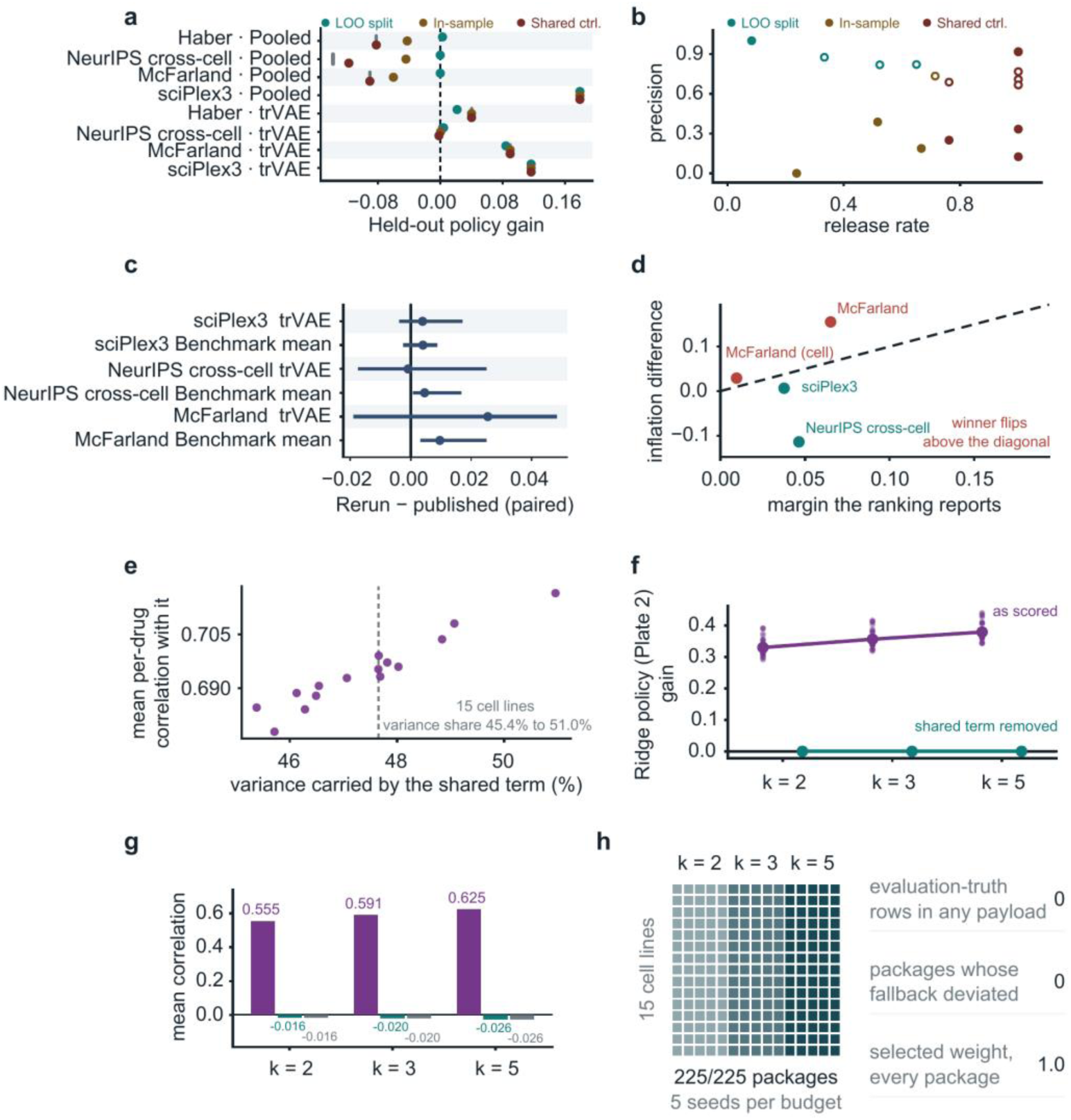
The criterion with a second base predictor, reproduction of published scores and blinded fits on the shallow plate. **a**, Gain after gating relative to not adapting for eight base-by-dataset rows (Pooled and trVAE as base, four datasets each) under three kinds of evidence: LOO split-control, in-sample split-control and shared-control evidence; grey tick, unconditional adaptation. **b**, Release rate against precision (fraction of released scored responses with positive gain) for the same rows; filled, Pooled as base; open, trVAE as base; two rows that released nothing are omitted. **c**, Paired per-operation difference between our re-run and the published score for six method-by-dataset pairs; point, median; bar, interquartile range. **d**, Difference in inflation between the top two methods against the margin reported under shared controls, for four settings; dashed line, the flip condition. **e**, Share of each context’s response variance carried by the single additive term common to all 88 drugs against the mean per-drug correlation with that term, one point per cell line (*n* = 15, shallow plate); dashed line, median share. **f**, Ridge policy (Plate 2) gain per trial at *k* = 2, 3 and 5 on the shallow plate as scored and after the shared term is removed from prediction and truth alike. **g**, The same removal on the calibration-mean arm (*n* = 900 splits) against the value obtained by centring alone on independent noise. **h**, Checks of the blinded Ridge policy (Plate 2) fits, which were fixed before held-out outcomes were read. Rows, 15 cell lines; columns, five seeds at each calibration budget (*k* = 2, 3 and 5), 225 fits in all. No fit read a held-out response, none departed from the recorded fallback, and every fit selected weight 1.0.

**Extended Data Fig. 7.**
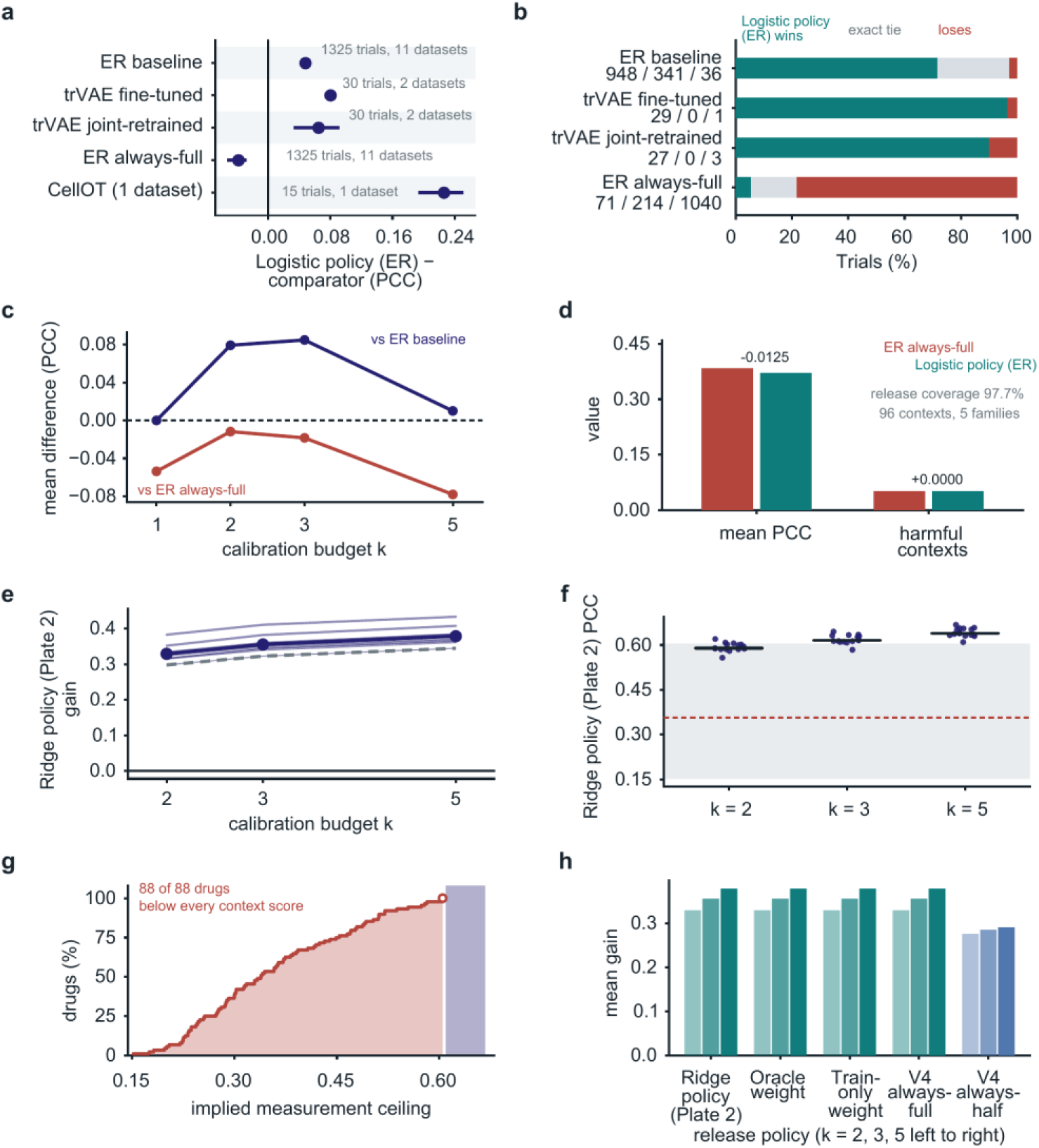
Gated adaptation compared with parameter updating, and measurement reliability on the shallow plate. a-d, Logistic policy (ER); e-h, Ridge policy (Plate 2). All panels use the shared-control scoring of their original comparisons and are not comparable with Fig. 3. **a**, Difference in PCC between Logistic policy (ER) and each comparator; bars, the run’s hierarchical interval where recorded and a context-clustered interval otherwise; the number of trials and datasets is given beside each row. Neural optimal transport (CellOT)^13^ was scored on one dataset. **b**, Trial-level outcome against each comparator (wins, exact ties, losses); counts beside each bar. **c**, Mean difference in PCC between Logistic policy (ER) and the ER baseline or ER always-full at calibration budgets of 1, 2, 3 and 5. **d**, Mean PCC and fraction of harmful contexts for ER always-full and Logistic policy (ER) on the 96 contexts of five external study families at *k* = 2. **e**, Gain per context under Ridge policy (Plate 2) on the shallow plate at *k* = 2, 3 and 5, one line per context; heavy line, median. **f**, Achieved correlation per context at three budgets against the band of per-drug ceilings implied by split-half reliability at 20 control cells; dashed line, mean ceiling; black bars, median context scores. **g**, Cumulative distribution of those ceilings over the 88 drugs; bar, lowest context score at *k* = 5. **h**, Mean gain of five release policies (Ridge policy (Plate 2), oracle weight, train-only weight, V4 always-full, V4 always-half) at *k* = 2, 3 and 5.

**Extended Data Fig. 8.**
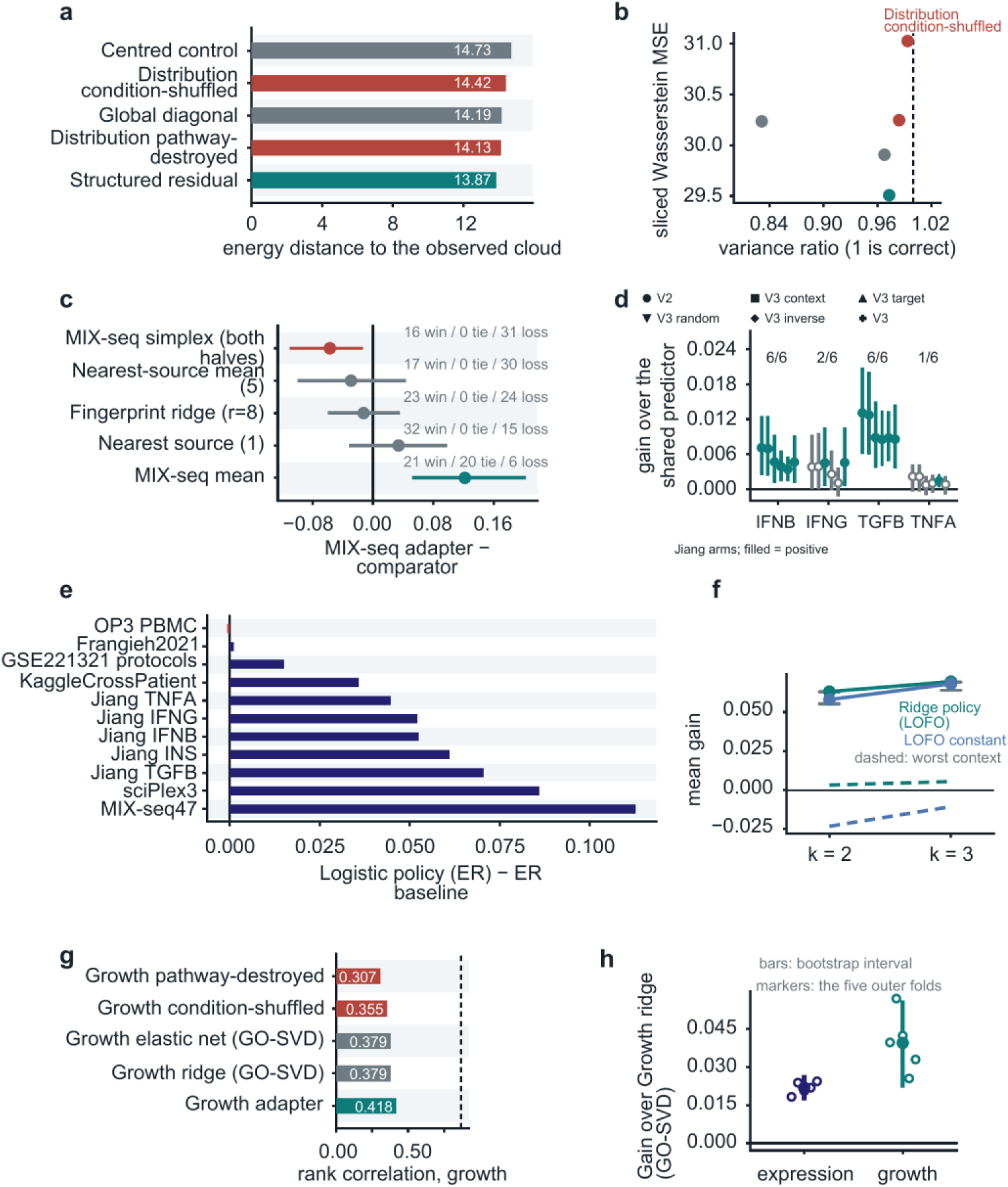
Distributional metrics, additional comparators, external study families and a growth phenotype. **a**, Energy distance^53^ from each of five arms to the observed cell cloud: structured residual, global diagonal, centred control, pathway structure destroyed, conditions shuffled. **b**, Sliced Wasserstein MSE against variance ratio for the same arms; dashed line, variance ratio of 1. **c**, Difference in PCC between the MIX-seq comparator-suite adapter and five comparators on 47 contexts and 282 context-target pairs; bars, hierarchical intervals of the comparator suite; counts, wins, ties and losses over the 47 contexts. **d**, Gain over the shared predictor for six adapter variants under four stimulus programmes (IFNB, IFNG, TGFB, TNFA); filled, strictly positive gain, with the number of positive variants printed above each programme; bars, 95% interval. **e**, Mean difference for Logistic policy (ER) relative to ER baseline for each of the eleven dataset families of the formal run. **f**, Mean gain of Ridge policy (LOFO) with one study family held out and of a leave-one-family-out constant, at *k* = 2 and 3 (930 trials, 93 contexts); dashed lines, worst contexts; grey lines, 97.5th percentile of the null. **g**, Spearman correlation between each arm’s predicted expression and a growth phenotype from a separate screen^54^ over 1,548 targets; dashed line, replicate reliability of the growth readout. **h**, Advantage over Growth ridge (GO-SVD) fold by fold on the expression and growth endpoints; bars, bootstrap interval; markers, the five outer held-out-target folds.

**Extended Data Fig. 9.**
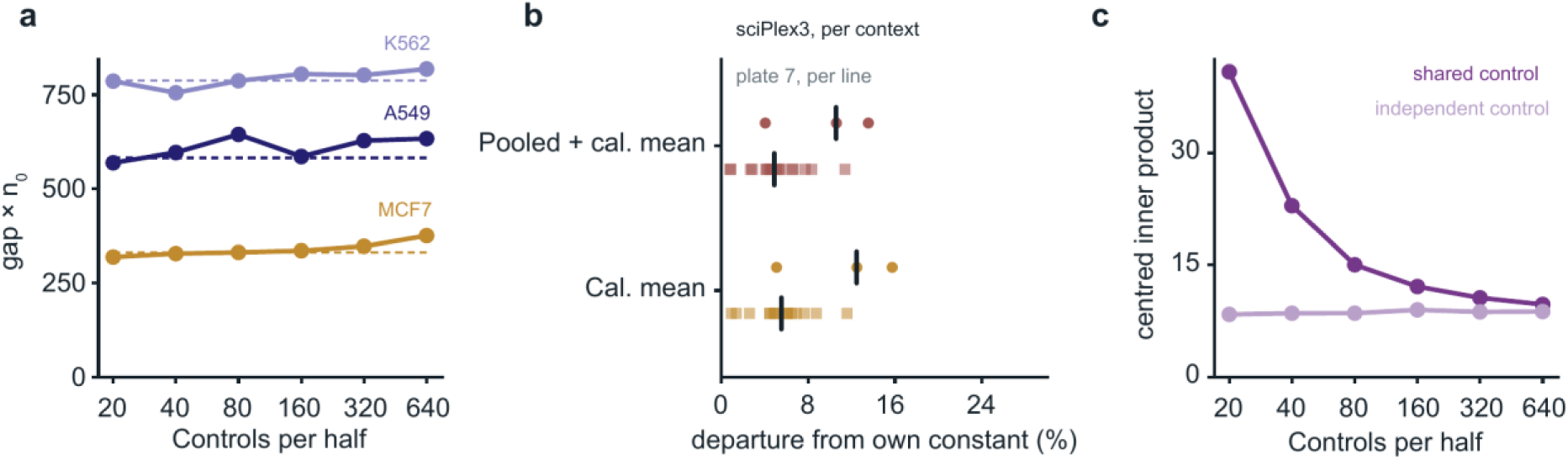
The depth invariant on a second dataset. sciPlex3, three contexts (K562, A549, MCF7), 33 drug-dose perturbations, plates as the control-split key, using the same nested-subsampling principle as the deep-plate depth sweep (Fig. 4e-g), but different treated-cell counts and depth levels: 504 scored responses per depth, 25 treated cells per condition held fixed, 20 to 640 control cells per half, three closed-form estimators. **a**, Product of the inner-product gap and control depth *n*_0_ (equation (11)) per context against control depth; dashed line, each context’s pooled median. **b**, Largest departure of a per-depth median from its own unit’s constant, one point per sciPlex3 context and per deep-plate cell line; bar, median; the control-free estimator is omitted because its constant is zero. **c**, Shared- and independent-control inner products for Pooled + cal. mean, against control depth per half.

