## Supplementary Note 1 for "A split-control criterion for few-shot adaptation in single-cell perturbation prediction"

### Supplementary Note 1 | Shared-control inflation: propositions, derivations and simulation checks

This note gives the statements and derivations behind equations (1)–(8) of the Online Methods, and the design and results of the simulations referred to there. Equation numbers without a prefix refer to the Online Methods; equations of this note are numbered (S1), (S2), ….

#### 1. Notation and assumptions

Fix one context. Let $d$ be the number of genes and index perturbations by $p$.

| Symbol | Meaning |
|---|---|
| $\mu_0$ | true mean profile of the unperturbed state |
| $\Delta_p$ | true response to perturbation $p$ |
| $T_p = \mu_0 + \Delta_p + \eta_p$ | measured pseudobulk profile under $p$; $\eta_p$ is the treated-cell sampling noise |
| $\hat{\mu}_0 = \mu_0 + \varepsilon_0$ | control estimate from $n_0$ control cells; $\varepsilon_0$ is its error |
| $D_p = T_p - \hat{\mu}_0 = \Delta_p + \eta_p - \varepsilon_0$ | observed response, equation (2) |
| $D'_p = \Delta_p + \eta_p - \varepsilon'_0$ | the same response referenced to an independent control estimate of the same depth |
| $\mathcal{K}, k = \lvert\mathcal{K}\rvert$ | calibration set and its size |
| $B_p$ | shared predictor, which reads no cell of the target context |
| $\sigma^2$ | per-gene variance of a single control cell |

All vectors are centred over genes. Writing $C = I - \mathbf{1}\mathbf{1}^\top/d$ for the centring projector, $\langle u, v\rangle = (Cu)^\top(Cv)$ and $\lVert u\rVert^2 = \langle u, u\rangle$. The Pearson correlation across genes is $\mathrm{PCC}(u, v) = \langle u, v\rangle/(\lVert u\rVert\lVert v\rVert)$.

Two scoring regimes are compared. Under shared controls the prediction and the observed response are referenced to the same control estimate, so both contain $\varepsilon_0$. Under independent controls the observed response is $D'_p$, with $\varepsilon'_0$ independent of $\varepsilon_0$ and identically distributed.

**Assumptions.**

- **A1 (additive, shared control error).** The control error enters every observed response of the context additively and as the same vector $\varepsilon_0$, with $\mathbb{E}\varepsilon_0 = 0$.
- **A2 (independent treated noise).** The $\eta_p$ are independent across perturbations and independent of $\varepsilon_0$ and $\varepsilon'_0$.
- **A3 (held-out target).** The scored perturbation is outside the calibration set, $p \notin \mathcal{K}$.
- **A4 (scalar carriage).** The prediction can be written as $P_p = P_p^0 - a\varepsilon_0$ with a constant $a$ and a component $P_p^0$ that is independent of $\varepsilon_0$ and $\varepsilon'_0$, equation (3).
- **A5 (equal depth).** $\varepsilon_0$ and $\varepsilon'_0$ are errors of control estimates with the same number of cells $n_0$.
- **A6 (isotropic control noise).** The entries of a control cell's noise are uncorrelated across genes with common variance $\sigma^2$, so that $\mathrm{Cov}(\varepsilon_0) = (\sigma^2/n_0)I$.

Under A6,

$$\mathbb{E}\|\varepsilon_0\|^2 = \mathrm{tr}(C\frac{\sigma^2}{n_0}I) = \frac{(d-1)\sigma^2}{n_0}, \qquad \text{(S1)}$$

because the centring projector has trace $d-1$. For a general covariance $\Sigma$ of $\varepsilon_0$ the right-hand side is $\mathrm{tr}(C\Sigma)$.

**Where the assumptions apply.** A1 holds when responses are differences of means on the analysis scale, as in the per-cell normalisation, log1p transformation and averaging used here. A4 holds exactly for the offset adapter and for the calibration mean (Section 2, Step 1). It does not hold as stated for a predictor that is a non-linear function of the control cells, which includes the published generative models; for these we report the projection coefficient of Section 3, whose identity holds for any predictor.

### 2. Proposition 1: the inner-product gap

**Proposition 1.** Under A1–A6, for every $p \notin \mathcal{K}$,

$$\mathbb{E}\langle P_p, D_p \rangle - \mathbb{E}\langle P_p, D'_p \rangle = \frac{a\sigma^2(d-1)}{n_0}. \tag{S2}$$

This is equation (4). The right-hand side does not depend on $k$ or on $P_p^0$, and it is zero when $a = 0$.

**Derivation.**

*Step 1: an additive few-shot correction satisfies A4 with a coefficient that does not decay with* $k$. Consider the offset adapter of equation (9) with weight $w$,

$$P_p = B_p + \frac{w}{k} \sum_{q \in \mathcal{K}} (D_q - B_q).$$

Substituting $D_q = \Delta_q + \eta_q - \varepsilon_0$,

$$P_p = \left[B_p + \frac{w}{k} \sum_{q \in \mathcal{K}} (\Delta_q + \eta_q - B_q)\right] - \frac{w}{k} \sum_{q \in \mathcal{K}} \varepsilon_0 = P_p^0 - w\varepsilon_0. \tag{S3}$$

The control error is the same vector for every calibration response, so its average over $\mathcal{K}$ is $\varepsilon_0$ itself and carries no factor $1/k$; the treated-cell noise $\eta_q$ does average down and is absorbed in $P_p^0$. By A2 and because $B$ reads no cell of the target context, $P_p^0$ depends only on $B$, $\Delta$ and $\{\eta_q\}_{q \in \mathcal{K}}$ and is independent of $\varepsilon_0$. Hence A4 holds with $a = w$; in particular $a = 1$ when the mean calibration residual, or the mean calibration response, is added at unit weight, and $a = 0$ for the shared predictor.

*Step 2: shared controls.* Expanding with A4 and equation (2),

$$\langle P_p, D_p \rangle = \langle P_p^0, \varDelta_p + \eta_p \rangle - \langle P_p^0, \varepsilon_0 \rangle - a\langle \varepsilon_0, \varDelta_p + \eta_p \rangle + a\langle \varepsilon_0, \varepsilon_0 \rangle.$$

The second term has expectation zero because $P_p^0$ is independent of $\varepsilon_0$ and $\mathbb{E}\varepsilon_0 = 0$. The third has expectation zero because $\varDelta_p$ is fixed, $\eta_p$ is independent of $\varepsilon_0$ (A2) and $\mathbb{E}\varepsilon_0 = 0$. By (S1) the fourth has expectation $a(d-1)\sigma^2/n_0$. Therefore

$$\mathbb{E}\langle P_p, D_p \rangle = \mathbb{E}\langle P_p^0, \varDelta_p + \eta_p \rangle + \frac{a(d-1)\sigma^2}{n_0}. \qquad \text{(S4)}$$

*Step 3: independent controls.* With $D_p' = \varDelta_p + \eta_p - \varepsilon_0'$ the same expansion gives the cross term $a\langle \varepsilon_0, \varepsilon_0' \rangle$ in place of $a\langle \varepsilon_0, \varepsilon_0 \rangle$; its expectation is zero because the two errors are independent with mean zero. The term $\langle P_p^0, \varepsilon_0' \rangle$ has expectation zero by A4. Hence

$$\mathbb{E}\langle P_p, D_p' \rangle = \mathbb{E}\langle P_p^0, \varDelta_p + \eta_p \rangle. \qquad \text{(S5)}$$

*Step 4.* Subtracting (S5) from (S4) gives (S2). ■

**Remarks.**

1. *Independence of $P_p^0$ and $\eta_p$.* Steps 2 and 3 do not use it; it is needed only for Proposition 2. It holds for $p \notin \mathcal{K}$ (A3), and it holds within each fold of a leave-one-out construction on the calibration set, where the scored response is excluded from the fit.

2. *Calibration budget.* The right-hand side of (S2) contains no $k$. This rests on $a$ being a constant. If the weight of the correction is itself fitted to the calibration responses, $a$ is a random variable that may depend on $k$, and (S2) holds with $a$ replaced by its value in each realisation only if that value is independent of $\varepsilon_0$. The absence of a factor $1/k$ is therefore derived for fixed weights; for fitted weights the dependence of the inflation on the calibration budget is reported as an observation (Fig. 2b).

3. *Linear carriage.* If $P_p = P_p^0 - M\varepsilon_0$ for a fixed matrix $M$, the same steps give an expected gap $\mathbb{E}\langle M\varepsilon_0, \varepsilon_0 \rangle = \mathrm{tr}(CMC\varSigma)$, which reduces to $\mathrm{tr}(M\varSigma)$ when $M$ and $\varSigma$ act on gene-

centred vectors, the convention of the Methods, and to (S2) when $M = aI$ and $\Sigma = (\sigma^2/n_0)I$.

4. *Relation to the fixed-prediction case.* When the prediction is a fixed vector $\tilde{\mu}$ scored as $\tilde{\mu} - \hat{\mu}_0$, A4 holds with $P^0 = \tilde{\mu} - \mu_0$ and $a = 1$, and (S2) gives the gap $\sigma^2(d-1)/n_0$ of the analysis that conditions on the prediction[20].

### 3. Proposition 1′: the two-fold identity

Let the control cells of the context be divided into two disjoint halves with mean profiles $c_A$ and $c_B$, and let the treated profiles $T_p$ be held fixed. Write $D_p^A = T_p - c_A$ and $D_p^B = T_p - c_B$, and let $P_p^A$ and $P_p^B$ be the predictions built with half A and with half B. Define

$$\mathrm{Gap}(0) = \langle P_p^A, D_p^A \rangle - \langle P_p^A, D_p^B \rangle, \quad \mathrm{Gap}(1) = \langle P_p^B, D_p^B \rangle - \langle P_p^B, D_p^A \rangle.$$

**Proposition 1′.** With $\Delta P = P_p^A - P_p^B$ and $\Delta c = c_A - c_B$,

$$\frac{1}{2}[\mathrm{Gap}(0) + \mathrm{Gap}(1)] = -\frac{1}{2}\langle \Delta P, \Delta c \rangle \tag{S6}$$

for any predictor. If $\Delta P = -a\Delta c$, the right-hand side equals $\frac{a}{2}||\Delta c||^2$, which is equation (5).

**Derivation.** Because $D_p^A - D_p^B = c_B - c_A = -\Delta c$, the treated profile cancels in each gap:

$$\mathrm{Gap}(0) = \langle P_p^A, D_p^A - D_p^B \rangle = -\langle P_p^A, \Delta c \rangle, \quad \mathrm{Gap}(1) = \langle P_p^B, D_p^B - D_p^A \rangle = \langle P_p^B, \Delta c \rangle.$$

Adding the two gives $-\langle P_p^A - P_p^B, \Delta c \rangle$, which is (S6). No expectation is taken. ■

**Consequences.**

1. *Projection coefficient.* Define $\hat{a} = -\langle \Delta P, \Delta c \rangle / ||\Delta c||^2$. Then (S6) reads $\frac{1}{2}[\mathrm{Gap}(0) + \mathrm{Gap}(1)] = \frac{\hat{a}}{2}||\Delta c||^2$ identically. With $\hat{a}$ estimated from the same two halves, equation (5) is therefore a sample identity for the unnormalised gap and not a test of the scalar-carriage model.

2. *Predictors that do not read the target controls.* If $P_p^A = P_p^B$, then $\mathrm{Gap}(0) = -\mathrm{Gap}(1)$ and the two-fold average is exactly zero in every realisation.

3. *Norm ratio.* The ratio $\|\varDelta P\|/\|\varDelta c\|$ equals $\hat{a}$ if and only if $\varDelta P = -a\varDelta c$ with $a \geq 0$. For a linear map, $\varDelta P = -M\varDelta c$, the two-fold average is the quadratic form $\frac{1}{2}\langle M\varDelta c, \varDelta c\rangle$, which the projection coefficient reproduces exactly and the norm ratio does not unless $M\varDelta c$ is parallel to $\varDelta c$.

4. *Consistency with Proposition 1.* For two halves of $n_0$ cells each under A6, $\mathbb{E}\|\varDelta c\|^2 = 2(d-1)\sigma^2/n_0$, so the expectation of $\frac{a}{2}\|\varDelta c\|^2$ is $a(d-1)\sigma^2/n_0$, in agreement with (S2).

### 4. Proposition 2: the correlation scale

Let

$$S = \mathbb{E}\|\varDelta_p + \eta_p\|^2, \quad \lambda = \frac{(d-1)\sigma^2/n_0}{S}, \quad r^2 = \frac{\mathbb{E}\|P_p^0\|^2}{S}, \quad \rho = \frac{\mathbb{E}\langle P_p^0, \varDelta_p + \eta_p\rangle}{\sqrt{\mathbb{E}\|P_p^0\|^2}\sqrt{S}},$$

so that $\lambda$ is the ratio of equation (1), and define the second-moment forms of the two scores,

$$\mathrm{PCC}_{\mathrm{shared}} = \frac{\mathbb{E}\langle P_p, D_p\rangle}{\sqrt{\mathbb{E}\|P_p\|^2\mathbb{E}\|D_p\|^2}}, \quad \mathrm{PCC}_{\mathrm{indep}} = \frac{\mathbb{E}\langle P_p, D_p'\rangle}{\sqrt{\mathbb{E}\|P_p\|^2\mathbb{E}\|D_p'\|^2}}.$$

**Proposition 2.** Under A1–A6,

$$\mathrm{PCC}_{\mathrm{shared}} = \frac{\rho r + a\lambda}{\sqrt{(r^2 + a^2\lambda)(1+\lambda)}}, \quad \mathrm{PCC}_{\mathrm{indep}} = \frac{\rho r}{\sqrt{(r^2 + a^2\lambda)(1+\lambda)}}, \qquad \text{(S7)}$$

and their difference is

$$\mathrm{inflation} = \frac{a\lambda}{\sqrt{(r^2 + a^2\lambda)(1+\lambda)}}. \qquad \text{(S8)}$$

These are equations (6) and (7).

**Derivation.** *Numerators.* By (S4) and (S5), $\mathbb{E}\langle P_p, D_p\rangle = \rho r S + a\lambda S$ and $\mathbb{E}\langle P_p, D_p'\rangle = \rho r S$, using $\mathbb{E}\langle P_p^0, \Delta_p + \eta_p\rangle = \rho r S$ from the definitions of $\rho$ and $r$.

*Denominators.* For the observed response, $\|D_p\|^2 = \|\Delta_p + \eta_p\|^2 - 2\langle \Delta_p + \eta_p, \varepsilon_0\rangle + \|\varepsilon_0\|^2$; the cross term has expectation zero (A2, $\mathbb{E}\varepsilon_0 = 0$) and (S1) gives $\mathbb{E}\|D_p\|^2 = S(1+\lambda)$. The same holds for $D_p'$ by A5. For the prediction, $\|P_p\|^2 = \|P_p^0\|^2 - 2a\langle P_p^0, \varepsilon_0\rangle + a^2\|\varepsilon_0\|^2$; the cross term has expectation zero (A4) and $\mathbb{E}\|P_p\|^2 = S(r^2 + a^2\lambda)$.

Substituting and cancelling $S$ gives (S7); subtracting gives (S8). ■

**Remarks.**

1. *Object of the statement.* (S7) and (S8) are exact for ratios of second moments. The expectation of the realised correlation, $\mathbb{E}[\mathrm{PCC}(P_p, D_p)]$, is the mean of a ratio and differs from the ratio of means; the size of the difference in simulation is reported in Section 6. No analytical bound on this difference is given here. The Methods state Proposition 2 for the second-moment ratio.

2. *Evaluation per perturbation.* $\lambda$, $r^2$ and $\rho$ carry the index $p$ through $S$ and $P_p^0$. Because (S8) is convex in $r^2$, evaluating it at parameters pooled over perturbations is not equivalent to evaluating it for each perturbation and averaging; the Methods use the latter.

3. *Special cases.* For $a = 1$ and $r = 1$, $\mathrm{PCC}_{\mathrm{shared}} = (\rho + \lambda)/(1 + \lambda)$, the limit obtained for a fixed prediction[20]. For $a = 0$ the inflation is zero for every $\lambda$.

4. *Role of $r$.* At fixed $a$ and $\lambda$ the inflation (S8) decreases as $r^2$ increases, whereas the inner-product gap (S2) does not depend on $P_p^0$. The inner-product scale therefore separates the carriage coefficient from the strength of the prediction, and the correlation scale does not.

### 5. The control-depth condition

Dividing (S2) by the response signal $S$ gives the signal-normalised inner-product gap,

$$\frac{\mathbb{E}\langle P_p, D_p\rangle - \mathbb{E}\langle P_p, D_p'\rangle}{S} = a\lambda = \frac{a(d-1)\sigma^2}{n_0 S}. \tag{S9}$$

Let $m$ be a bias target on this scale. Requiring the gap not to exceed a quarter of the target, $a\lambda \le m/4$, and solving (S9) for $n_0$ gives

$$n_0 \ge \frac{4a(d-1)\sigma^2}{mS}, \tag{S10}$$

which is equation (8). The right-hand side is linear in $a$ and in the per-gene control variance and inverse in the response signal, so that at a given depth a context with weak responses requires more control cells for the same target. In the benchmark projections $m$ was set from the margin between adjacent methods, and the factor 4 expresses the target of a quarter of that margin; a target of a tenth replaces 4 by 10.

Equation (S10) concerns the inner-product scale. On the correlation scale, (S8) is not proportional to $1/n_0$, because $\lambda$ also enters its denominator. The depth projections of Fig. 2g hold that denominator constant and scale the observed PCC inflation by the ratio of depths; they are approximations and do not follow from (S10).

### 6. Simulation checks

#### 6.1 Proposition 2 in twelve settings

**Design.** Data were generated under A1–A6 with $d = 300$ genes and a per-cell control variance $\sigma^2 = 1.7$. For each of six control depths, $n_0 \in \{10,25,50,100,200,400\}$, and two carriage coefficients, $a \in \{1,2\}$, a true response $\Delta \sim \mathcal{N}(0, I_d)$ was drawn once and 30,000 replicates were simulated. In each replicate the two control errors $\varepsilon_0$ and $\varepsilon_0'$ were drawn independently from $\mathcal{N}(0, (\sigma^2/n_0) I_d)$, the treated-cell noise of the scored response and of five calibration responses from $\mathcal{N}(0, 0.3^2 I_d)$, and a control-free predictor as $B = 0.6\Delta + 0.4z$ with $z \sim \mathcal{N}(0, I_d)$. The prediction was $P = P^0 - a\varepsilon_0$ with $P^0 = B + a \cdot \frac{1}{5}\sum_q (\Delta + \eta_q - B)$. The realised noise-to-signal ratio $\lambda$ ranged from 0.0034 to 0.180 across the twelve settings, a 52-fold span, and $r^2$ was 0.92–0.94 for $a = 1$ and 2.01 for $a = 2$.

Two quantities were compared with the closed form (S8) evaluated at the realised $\lambda$ and $r^2$: the difference of the second-moment ratios of Section 4, with a standard error from the delta method, and the difference between the mean realised correlations under shared and under independent controls.

**Results.**

| $n_0$ | $a$ | $\lambda$ | $r^2$ | **closed form (S8)** | **second-moment ratio** | **residual** | $z$ | **mean of realised PCC** | **residual** | $z$ |
|---|---|---|---|---|---|---|---|---|---|---|
| 10 | 1 | 0.18039 | 0.9235 | 0.15803 | 0.15801 | −0.00002 | −0.18 | 0.15767 | −0.00036 | −3.87 |
| 10 | 2 | 0.15352 | 2.0103 | 0.17647 | 0.17648 | +0.00000 | +0.04 | 0.17614 | −0.00033 | −3.22 |
| 25 | 1 | 0.05852 | 0.9377 | 0.05699 | 0.05693 | −0.00006 | −1.33 | 0.05677 | −0.00022 | −5.12 |
| 25 | 2 | 0.06181 | 2.0101 | 0.07985 | 0.07992 | +0.00007 | +1.24 | 0.07972 | −0.00013 | −2.16 |
| 50 | 1 | 0.03241 | 0.9311 | 0.03250 | 0.03245 | −0.00005 | −1.49 | 0.03236 | −0.00014 | −4.67 |
| 50 | 2 | 0.02926 | 2.0078 | 0.03957 | 0.03959 | +0.00001 | +0.31 | 0.03948 | −0.00010 | −2.69 |
| 100 | 1 | 0.01490 | 0.9368 | 0.01517 | 0.01516 | −0.00001 | −0.39 | 0.01511 | −0.00006 | −2.94 |
| 100 | 2 | 0.01455 | 2.0076 | 0.02011 | 0.02013 | +0.00002 | +0.89 | 0.02007 | −0.00004 | −1.45 |
| 200 | 1 | 0.00726 | 0.9386 | 0.00743 | 0.00744 | +0.00001 | +0.79 | 0.00742 | −0.00001 | −1.13 |
| 200 | 2 | 0.00751 | 2.0097 | 0.01047 | 0.01046 | −0.00001 | −0.74 | 0.01043 | −0.00004 | −2.50 |
| 400 | 1 | 0.00344 | 0.9413 | 0.00354 | 0.00355 | +0.00001 | +1.18 | 0.00353 | −0.00000 | −0.10 |
| 400 | 2 | 0.00372 | 2.0083 | 0.00522 | 0.00523 | +0.00001 | +0.84 | 0.00521 | −0.00000 | −0.38 |

Against the second-moment ratio, the closed form was within three standard errors in all 12 settings; the largest absolute residual was 0.00007 (at most 0.28% of the inflation) and the residuals were of both signs. Against the mean of the realised correlations, the closed form was within three standard errors in 8 of 12 settings; the largest absolute residual was 0.00036 (at most 0.44% of the inflation) and all twelve residuals were negative, with the largest standardised residual at $-5.1$. Equation (S8) therefore describes the second-moment ratio to within Monte Carlo error over this range, and exceeds the inflation of the mean realised correlation by less than half a per cent of its value.

#### 6.2 Propositions 1 and 1′

Additional simulations under the same assumptions ($d = 300$, $n_0 = 50$, 40,000 replicates unless stated) gave the following.

| Check | Expected | Observed |
|---|---|---|
| Gap (S2), $a = 0$ | 0 | −0.0229 ($z = -1.38$) |
| Gap (S2), $a = 1$ | 10.166 | 10.1904 ($z = +1.10$) |
| Gap (S2), $a = 2$ | 20.332 | 20.3569 ($z = +0.70$) |
| Gap at $k = 2$, 5, 20 and 80 ($a = 1$) | constant in $k$ | 10.155, 10.160, 10.140 and 10.181; range 0.041, below three standard errors (0.072) |
| Gap as the correlation of $P^0$ with the response varied from 0.519 to 0.990 | constant | 10.117 to 10.175 |
| Gap $\times$ $n_0$ over an eight-fold range of $n_0$ | constant | 508, 507, 509 and 507; coefficient of variation 0.0017 |
| $\langle P^0, \eta_p \rangle$ under leave-one-out calibration | 0 | −0.0183 ($z = -0.54$) |
| Two-fold identity (S6), 300 draws at $a = 0$, 1 and 2.3 | exact | largest relative deviation $9.5 \times 10^{-16}$ |
| Linear carriage with a rank-16 component: projection coefficient in (S6) | exact | largest relative deviation $8.8 \times 10^{-16}$ |
| Same, norm ratio in place of the projection coefficient | not exact | median relative error 1.8% |

Each algebraic step of the derivations in Sections 2–4 was also checked symbolically; the simplified difference between the two sides was zero in every case.
